# Cortical dynamics underlying social behavior in dominance hierarchy and spatial navigation

**DOI:** 10.1101/2020.06.12.147249

**Authors:** Ariel Lara-Vasquez, Nelson Espinosa, Cristian Morales, Constanza Moran, Pablo Billeke, Joseph Gallagher, Joshua J. Strohl, Patricio T. Huerta, Pablo Fuentealba

**Affiliations:** Centro Integrativo de Neurociencias y Departamento de Psiquiatría, Pontificia Universidad Católica de Chile, Santiago, Chile; Centro de Investigación en Nanotecnología y Materiales Avanzados - CIEN-UC, Pontificia Universidad Católica de Chile, Santiago, Chile; Laboratorio de Neurociencia Social y Neuromodulación, Centro de Investigación en Complejidad Social, Universidad del Desarrollo, Santiago, Chile; Laboratory of Immune & Neural Networks, Feinstein Institutes for Medical Research, Manhasset, New York, USA; Department of Molecular Medicine, Zucker School of Medicine at Hofstra/Northwell, Manhasset, New York, USA

**Keywords:** Dominance hierarchy, Spatial navigation, Prefrontal cortex, Hippocampus, Cortical oscillations

## Abstract

Rodents establish dominance hierarchy as a social ranking system in which one subject acts as dominant over all the other subordinate individuals. Dominance hierarchy regulates food access and mating opportunities, but little is known of its significance in collective behavior, for instance during navigation for foraging or migration. Here, we implemented a simplified goal-directed spatial navigation task in mice and found that the social context exerts significant influence on individual decision-making, even when efficient navigation rules leading to reward had been previously learned. Thus, decision-making and consequent task performance were strongly dependent on contingent social interactions arising during collective navigation, yet their influence on individual behavior was outlined by dominance hierarchy. Dominant animals did not behave as leaders during navigation; conversely, they were most sensitive to social context. Social ranking in turn was reflected in the neural activity and connectivity patterns of the prefrontal cortex and hippocampus, both in anesthetized and behaving mice. These results suggest that the interplay between contingent social interactions and dominance hierarchy can regulate behavioral performance, supported by the intrinsic matrix of coordinated activity in the hippocampal-prefrontal circuit.

**Significance Statement:** Decision-making is shaped by intrinsic features, such as memory-stored information, and external influences, such as social interactions, yet their interplay is not well understood. We studied decision-making during collective behavior and found that instead of prioritizing memory-based pertinent information, mice shifted their individual decisions according to contingent social interactions arising in the social context. Conversely, constitutive social interactions, such as dominance hierarchy, were relevant to outline the effect of the social environment on individual behavior. Our results suggest that intrinsic hippocampal-cortical activity and connectivity patterns define social interactions. Hence, intrinsic cortical dynamics underlie behavioral performance during social decision-making.

## Introduction

Social behavior is an adaptive response that has evolved to improve ecological fitness in many species [1]. Mammalian social behaviors occur in the context of extended groups; however, in laboratory settings, interactions such as fighting, chasing, courtship, and grooming, are typically investigated in pairs of individuals [2, 3]. This approach of studying dyads, and treating the results as prototypical social behavior, has significant limitations because animal groups commonly rely on more complicated social structures. Indeed, recent experiments tracking mice in ethologically relevant environments have revealed strongly correlated social behaviors that become evident in settings of not just two but multiple individuals [2, 3]. Moreover, some social behaviors that arise in groups can be contingent, as occurs when individuals randomly meet during environmental exploration, or when they court a mating partner [4]. Conversely, social interactions can be a constitutive group property, such as the social ranking system [5]. Dominance status in a social group can be important as it regulates individual behavior in assays evaluating anxiety, locomotion, or aggressiveness, to an extent comparable to genetic mutations or pharmacological agents [6, 7]. Nevertheless, most reports describe the role of dominance status only under individual conditions.

Previous studies have established that the neural basis of dominance hierarchy relies on the efficacy of synaptic transmission in the medial prefrontal cortex (mPFC) [8], particularly that of pyramidal cells [9]. Furthermore, synaptic activity in the mPFC as well as its functional connectivity with the dorsal hippocampus are essential to regulate spatial navigation and decision-making [10]. More generally, it has been proposed that the mPFC processes the current context and compares it with past experience to predict and execute the most adaptive behavioral response [11]. Indeed, the mPFC is actively recruited and prominently contributes to the control of dominance hierarchy [9], spatial navigation [12], and social behavior [13]. In these cases, both activity and connectivity of the mPFC have been dynamically associated with ongoing behavioral needs. However, it is not known whether the dynamics of the mPFC network accounts for performance of executive function in social contexts. Here, we tested the hypothesis that the connectivity matrix of the mPFC correlates with specific features of social behavior and dominance hierarchy. We developed a simple social navigation task and performed cortical recordings, in both anesthetized mice and freely-moving mice. Our results show that dominance hierarchy is represented in the activity and connectivity patterns of the mPFC and hippocampus, which also modulate social interactions during goal-directed spatial navigation.

## Results

### Social interactions affect decision-making during goal-directed spatial navigation

We first characterized the influence of social context on individual decision-making during collective spatial navigation. We exposed groups of 4 littermate males to a spatial navigation task, based on a modified version of the T-maze [14], which accommodated all mice simultaneously (**Fig. 1A**). As a result, our T-maze was larger than standard versions (Fig. S1). We structured the task in 2 temporally ordered phases consisting of a training phase (10 trials per day for each mouse) followed by a testing phase. During training, 2 littermates were pseudo-randomly assigned to individually look for reward exclusively in the left arm, while the right arm was baited for the remaining 2 littermates. There was large variance in performance (defined as the proportion of correct choices during individual trials) between individuals during the training phase (Fig. S2), but overall, mice improved their performance linearly over time (Fig. S3). In addition, they progressively decreased their latency, defined as the time interval taken to reach the rewarded pocket in the baited arm (Fig. S3). Throughout sessions, performance and latency co-varied linearly (Fig. S3) suggesting interdependence (Table S1). Littermates were sequentially trained on the same session, and since learning rates varied among mice, we defined a learning criterion based on litter performance (3 of 4 mice with 0.75 performance during 2 consecutive days).

**Figure 1.**
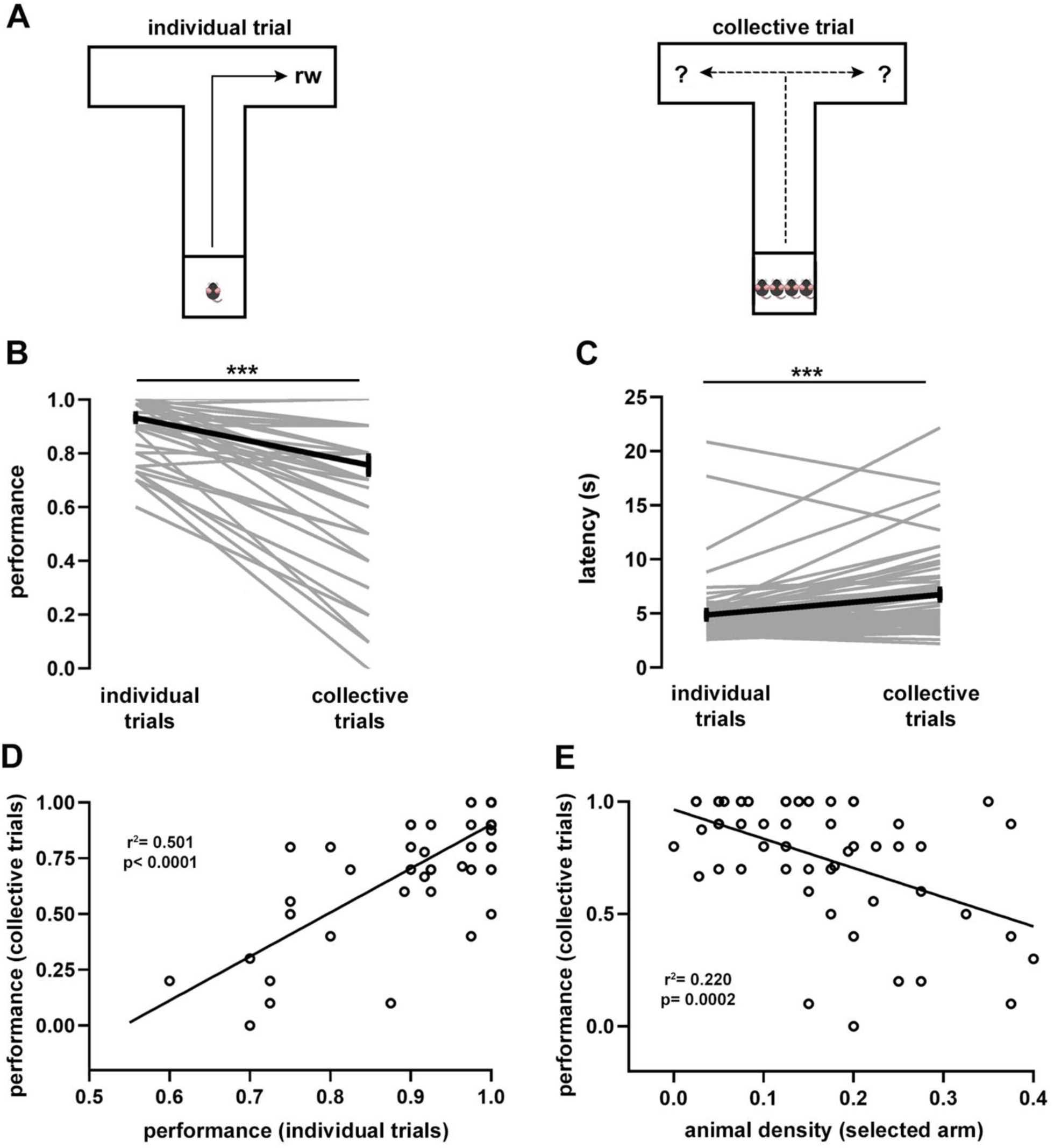
Spatial navigation task and associated behavioral performance. A, T-maze spatial navigation task in two variants. Littermate mice (n = 60) were individually trained (10 trials per day) to navigate the maze foraging for food located at the end of an arm until reaching the learning criterion. Thereafter, four individual trials (reward in fixed location) were alternated with one collective trial (reward in random location). From every litter, two randomly chosen mice were consistently trained to look for food in one arm and the remaining two mice in the opposite arm. Average task performance (B) and latency (C) for individual mice during collective and individual trials sampled during the testing phase. D, Spearman correlation between the average performance of individual trials against the average performance of collective trials. R^2^ = 0.501, P < 0.0001. E, Spearman correlation between the proportion of mice located in the selected arm against average navigation performance of individual mice during collective trials. R^2^ = 0.22, P = 0.002. Wilcoxon signed rank test (***, P < 10e-6). Gray lines, individual mice; black lines, population averages ± SEM.

Once the learning criterion was reached, mice started the testing phase, which consisted of 4 consecutive individual trials that were followed by a collective trial, in which all 4 littermates were tested simultaneously (**Fig. 1A**). To prevent learning of the reward location during collective trials, arms were randomly baited (in every collective trial). We found that, during the testing phase, both performance and latency of individual trials reached a plateau and were stable over the remaining testing period, suggesting that the task had been acquired and consolidated (Fig. S3). Notably, comparison of performance during individual (0.93 ± 0.01) and collective (0.75 ± 0.03) trials showed a significant drop in task performance (P = 2.72e-08, **Fig. 1B**), suggesting an important effect of social context in the process of decision-making of individual mice during collective navigation. The performance drop in collective trials was largely determined by task acquisition in the previous training phase, as they were strongly correlated (P = 1.5e-10, **Fig. 1D**). Conversely, task latency during the testing phase increased when comparing individual (4.99 ± 0.40 s) and collective (6.86 ± 0.49 s) trials (P = 4.86e-07, **Fig. 1C**). Increased latency was proportional to the previous task acquisition since mice exhibiting short latency during individual trials increased less their latency during collective trials (P = 4.4e-11, Fig. S3). Hence, the social context produced a shift in decision-making, which was reflected in performance decay and a proportional latency increase during goal-directed, collective spatial navigation.

Movement decisions in animal groups often depend on contingent social interactions among individual subjects [4, 15]. During collective movement, animals tend to be attracted to conspecifics to avoid being isolated and to align themselves with neighbors [16, 17]. Thus, we reasoned that, during collective movement, mice might modify their previously learned trajectory depending on the distribution of animals in the maze arms. To test this idea, we calculated for every mouse the relative average density of animals located in the selected arm and projected it against its average performance during collective trials (P = 0.0002, **Fig. 1E**). We found that, during collective trials, performance was inversely proportional to the relative density of animals in the selected arm. Conversely, there was no relation between the proportion of animals located in the opposite arm and performance during collective trials (P = 0.293, Fig. S4). This implies that the density of animals in the arm that a given mouse chose to move into was correlated with its task performance in the social context. To further explore this observation, we used a mixed logistic model to assess the influence of the spatial distribution of animals on task performance during collective navigation (Tables S2 and S3). We confirmed that animal density in the arms exerted significant influence in shifting the decision-making strategy during collective navigation, with particular relevance to the proportion of mice located in the selected arm (P = 6.41e-11). Therefore, when more mice accumulated in a lateral arm it was more likely that an individual mouse would move to that arm, regardless of the previously learned reward location. Thus, contingent social interactions were able to modulate memory-based learned reward values and bias decision-making during collective spatial navigation.

### Dominance hierarchy outlines the influence of social context during spatial navigation

We explored the role of dominance hierarchy [5] on social interactions during spatial navigation. We assessed hierarchical relations of mice with the tube test [9] in parallel to the spatial navigation task described above. This test measures the dominance tendency by placing pairs of mice in a narrow tube, facing each other, and one mouse forces the other out backward to obtain victory (**Fig. 2A**). We established social ranking based on the success rate of mice in pair-wise tests, using a round robin design (**Fig. 2A**), and found that interaction time in the tube was shorter as ranking difference increased (P = 2.31e-13, **Fig. 2B**). Dominance hierarchy was stable over time, particularly for the dominant mouse, whose position was rarely challenged throughout the experimental protocol (Fig. S5). Interestingly, the dominant mouse was not the largest animal in the group, as body masses were similar between rankings during both ad libitum access to food (P = 0.8216, Fig. S6) and food restriction periods in the navigation test (P = 0.6554, Fig. S6). During the training phase, we observed that social ranking was not relevant for task acquisition as performance and latency were comparable between animals regardless of their dominance hierarchy (Fig. S3). Similarly, the time required to reach the learning criterion was not different between social rankings (P = 0.9609, Fig. S7). During the testing phase, the performance drop and latency increase in collective trials was not modulated by dominance hierarchy, as it was not different between social groups (performance, P = 0.6552; latency, P= 0.5662; Fig. S7).

**Figure 2.**
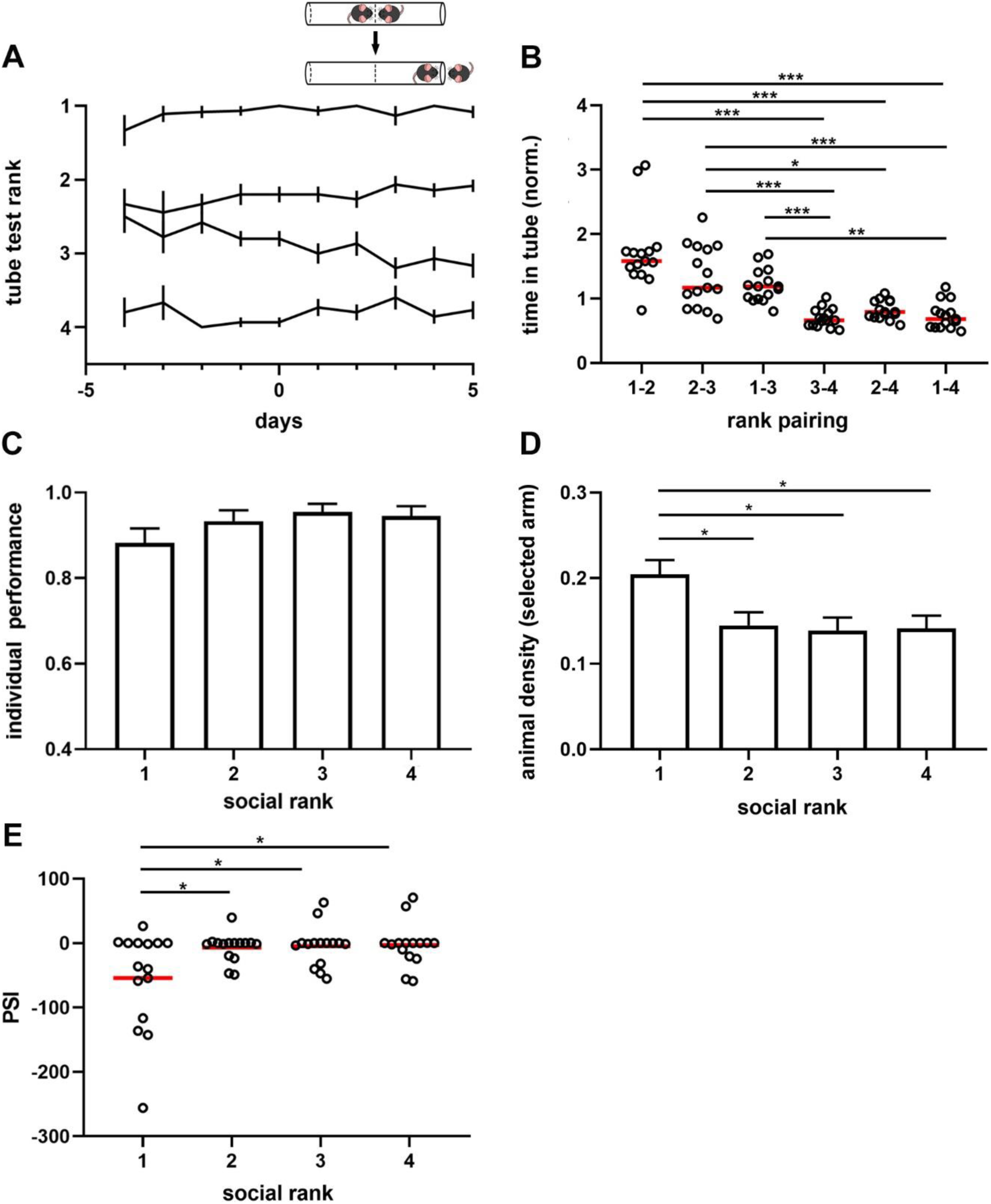
Dominance hierarchy and sensitivity to social context during spatial navigation. A, summary plot for all measured cages (n = 15). Lines show average rank position based on the proportion of victories in the tube test (inset) correlative to the spatial navigation test (days - 4 to 0, training phase; days 1 to 5, testing phase). Ranking 1, dominant; ranking 2, first active subordinate; ranking 3, second active subordinate; ranking 4, submissive. Note ranking stability over time, particularly for dominant mice. Inset, schematic of the tube test used to identify the mice ranking system. B, normalized time spent in the tube for the six pairing conditions. One-way ANOVA, P = 2.31e-13. C, average performance of individual trials during the testing phase by social ranking. One-way ANOVA, P = 0.25. D, animal density in the arm selected by animals during the navigation task according to social ranking. One-way ANOVA, P = 0.007. E, peer sensitivity index (PSI) by social ranking. One-way ANOVA, P = 0.0084. Bonferroni test post hoc (*, P < 0.05; **, P < 0.01; ***, P < 0.001). Black lines, population averages ± SEM; circles, individual mice average; red line; population average; bars represent average ± SEM.

Next, we assessed the influence of dominance hierarchy on task acquisition and found no significant difference of individual performance across social ranks in the navigation task (P = 0.25, **Fig. 2C**). We then evaluated the effect of social contingent interactions arising during navigation, as this is another factor that might modulate performance. We compared animal density in the selected arm across social ranks during collective navigation and found that, when compared to the subordinate groups, dominant animals moved more eagerly to the arm that was more densely populated, regardless of the reward location (P = 0.007, **Fig. 2D**). No such difference was detected in the opposite arm (P = 0.4499, Fig. S4). This result suggested that dominant mice may be more influenceable by the social context than other groups. We reasoned that the predisposition of individual mice to change their decision according to the distribution of littermates in the maze should be related to the influence of social context on individual behavior. To obtain an estimate of social influence, we computed for every individual mouse the regression coefficient of the spatial distribution of mice on the maze against task performance in the social context and called it the ‘peer susceptibility index’ (PSI, median = 0, IQR = 35.13). Since PSI was proportional to the social influence on individual behavior, the larger its value, the stronger the effect of social context on task performance. Thus, negative values reflect a detrimental effect of social context, whereas positive values indicate a beneficial effect on task performance. Importantly, PSI was significantly different between dominant mice and subordinate groups (P = 0.0084, **Fig. 2E**), thus suggesting that dominant mice were more likely to shift their decision based on social context. Moreover, differences in PSI did not result from different overall distributions of littermates in the maze during collective navigation according to social ranking (P = 0.1783, Fig. S4). Altogether, these results suggest that mice exhibit differential susceptibility to contingent social interactions, outlined by dominance hierarchy.

### Activity and connectivity in the hippocampal-prefrontal circuit correlate with behavioral performance

Previous studies have established the neural basis of dominance hierarchy in the synaptic connectivity of the mPFC [9, 18], hence we studied the relation between intrinsic mPFC dynamics and social ranking system. We recorded spontaneous rhythmic cortical activity in animals with stereotaxically-implanted electrodes in mPFC and hippocampus (Fig. S8), two brain regions that are required for goal-directed spatial behavior [19]. Initially, we performed the experiments under deep anesthesia to focus on intrinsic cortical dynamics and to minimize behavioral confounds resulting from different dominance hierarchy. To compare relatively similar conditions, we assessed the depth of anesthesia, as revealed by the power of the delta frequency band (0.5–4 Hz) of the mPFC and found no differences between social ranks (P = 0.081, Fig. S9). This result suggested that the global brain state was roughly similar across social groups.

Analysis of the hippocampal oscillations showed that all animals exhibited epochs of prominent sharp wave-ripples (SWRs, 100–250 Hz, **Fig. 3**) that alternated with spontaneous theta-band oscillations (4–8 Hz, **Fig. 4**), characteristic of hippocampal quiescent [20] and exploratory [21] states, respectively. We first probed the quiescent states of the brain network (**Fig. 3**), when SWRs dominate hippocampal activity [22], and found prominent SWR events (**Fig. 3A**), which occurred as short-lived, fast waxing and waning oscillations (**Fig. 3B**). Remarkably, SWR amplitude was dependent on social ranking as dominant mice exhibited the largest SWRs when compared to the subordinate groups (P = 8.55e-18, **Fig. 3C**). The duration of SWRs was not different between social ranks (P = 0.0658), yet their frequency was slower in the submissive group (P = 5.45e-38, **Fig. 3D**).

**Figure 3.**
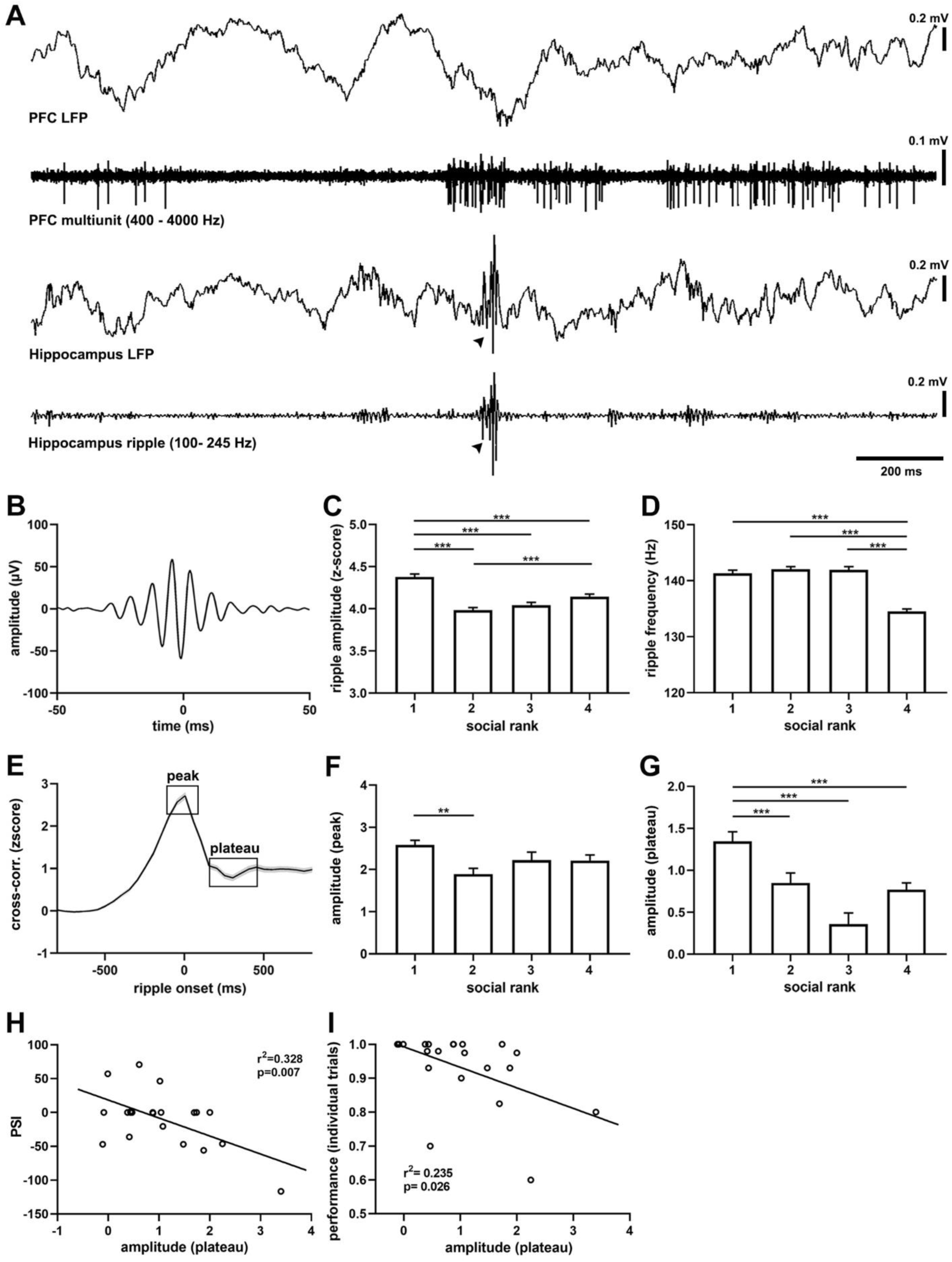
Sharp wave ripples in cortical networks across social groups. A, example simultaneous recordings of hippocampal (LFP HP) and prefrontal cortical (LFP PFC) activity showing hippocampal sharp wave ripples (SWRs, filtered 100-250 Hz) and cortical spiking activity (units PFC, filtered 300-4000 Hz) recorded in a urethane-anesthetised mouse (CM99reg05). B, grand average ripple episode (n = 18,571 events, 21 animals). Peak SWRs amplitude (C) and frequency (D) by social ranking. One-way ANOVA, C, P = 8.55e-18; D, P = 5.45e-38. E, average crosscorrelogram between the onset of SWRs and PFC units (n = 658 units). Note sustained component (plateau) after maximal activity (peak). F, crosscorrelogram peak amplitude by social ranking. One-way ANOVA, P = 0.002. G, crosscorrelogram plateau amplitude by social ranking. One-way ANOVA, P = 2.81e-07. H, linear regression between plateau amplitude against peer sensitivity index (PSI). R^2^ = 0.328, P = 0.007. I, linear regression between plateau amplitude against average performance during individual trials. R^2^ = 0.235, P = 0.026. Bonferroni test post hoc (**, P < 0.01; ***, P < 0.001). Black lines, population averages; shading areas, ± SEM; bars, average ± SEM.

**Figure 4.**
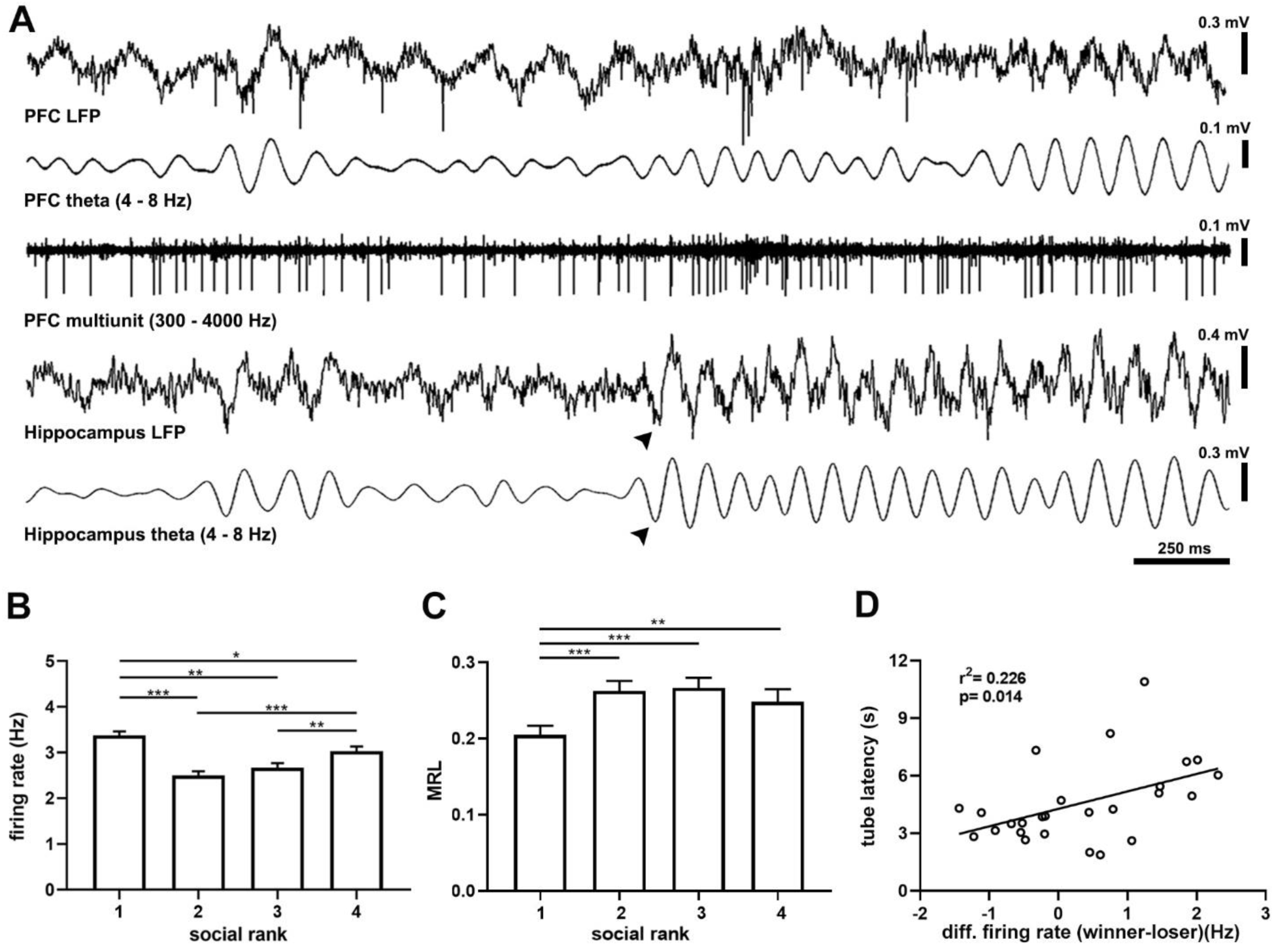
Theta oscillations in cortical networks across social groups. A, example simultaneous recordings of hippocampal (LFP HP) and prefrontal cortical (LFP PFC) activity showing theta oscillations (filtered 4-8 Hz) and cortical spiking activity (units PFC, filtered 300-4000 Hz) recorded in a urethane-anesthetized mouse (CM24reg05). Arrowhead depicts the onset of theta oscillations. B, average prefrontal single-unit firing rate by social ranking (n = 3,702 units). One-way ANOVA, P = 5.56e-11. C, mean resultant length (MRL) of the population vector from single-unit activity in relation to cortical theta oscillations according to social raking. Kruskal-Wallis test, P = 3.10e-06. D, linear regression between prefrontal cortex firing rate difference (winner – loser) and tube test latency difference (winner – loser). R^2^ = 0.226, P = 0.014. Bonferroni test post hoc (*, P < 0.05; **, P < 0.01; ***, P < 0.001). Bars, average ± SEM.

SWRs powerfully synchronize neuronal spike-timing across the neocortex [23, 24], including the mPFC [25, 26]. Therefore, we computed cross-correlation functions between hippocampal SWRs and mPFC spikes to quantitate the degree of synchronization between them. We found that cortical spiking activity increased preceding the onset of hippocampal SWRs (**Fig. 3E**), yet maximal mPFC activation (when SWRs peaked) exhibited little difference between social groups (**Fig. 3F**). After SWR episodes, cortical activation decreased rapidly but did not return immediately to baseline. Instead, it reached a plateau of sustained neuronal discharge (**Fig. 3E**). In paired hippocampal-cortical recordings, this plateau activation was evident as a prolonged after-discharge of mPFC neurons well beyond the end of the SWR (Fig. S10). Notably, plateau post-ripple discharges were stronger in dominant mice when compared to subordinate groups, suggesting enhanced functional connectivity following SWR episodes (P = 2.81e-07, **Fig. 3G**). These observations were robust, confirmed by shuffling comparisons, and apparent when assessing the entire neuronal population (P = 7.71e -47, Fig. S11). Moreover, the differences in plateau discharges were not the result of different temporal distributions of SWRs, as inter-ripple intervals were similar across social groups (P = 0.6193, Fig. S11).

Our data showed that both cortical connectivity and susceptibility to social context were dependent on social ranking, particularly for dominant mice. We thus reasoned that the PSI, a behavioral parameter, might be correlated with the plateau discharge, a neurophysiological parameter. Indeed, the PSI significantly correlated with the plateau discharge (P = 0.00666, **Fig. 3H**), but not with the peak cross-correlogram (Tables S4 and S5). We then tested whether intrinsic hippocampal-cortical connectivity was related to task performance during spatial navigation. We performed multiple linear regressions between behavior (task performance) and neural activity (SWRs) and found that the amplitude of the plateau discharge correlated with both performance (P = 0.026, **Fig. 3I**) and latency (P = 0.0019, Fig. S11) of individual trials, but not with collective trials (Tables S6 and S7). Since the PSI reflects the influence of social context on task performance, these results suggest that the strength of intrinsic cortical connectivity correlates with animal behavior during spatial navigation.

We then assessed activated cortical states (**Fig. 4**) that were characterized by prominent theta oscillations [27]. The spectral distribution of field potentials evidenced strong hippocampal theta that was consistently similar between social groups, with comparable amplitude, frequency, and duration (Fig. S12). Theta waves were also detected in mPFC and were associated with hippocampal theta (**Fig. 4A**). Since oscillatory synchrony is a neural mechanism for functional coupling of distributed neural circuits [28], we assessed the spontaneous spectral coherence in the field potential activity of the hippocampal-cortical circuit. We identified elevated intercortical coherence in theta oscillations under anesthesia in all animals, with no difference between social rankings (Fig. S12). In vitro studies in mPFC slices have shown that dominant animals exhibit larger synaptic strength in their excitatory synapses than submissive mice [9]. We thus tested whether this in vitro relation translated to in vivo spiking patterns and found that the overall mPFC firing rate in dominant mice was larger than the subordinate groups (P = 5.56e-11, **Fig. 4B**, Table S8). Interestingly, the difference was specific to regular spiking cells (P = 6.07e-12, Fig. S13), which are putative pyramidal neurons, as it was not detected in fast spiking units (P = 0.6243, Fig. S13), which are putative interneurons. Hence, dominant mice exhibited larger levels of intrinsic spiking activity in the mPFC under anesthesia. Cortical oscillations synchronize neuronal spiking and contribute to the temporal integration of neural activity. Therefore, we computed the oscillatory phase-locking of mPFC neurons to hippocampal theta by calculating the mean resultant length (MRL) of individual spikes to theta cycles [26]. As a population, mPFC neurons from dominant animals were less strongly modulated by hippocampal theta than subordinate mice (P = 0.0026, Fig. S12). Similarly, mPFC neurons from dominant animals were less modulated by local cortical theta than cells from the subordinate groups (P = 3.10e-06, **Fig. 4C**).

Recent evidence shows that the firing rate of mPFC neurons correlates with effortful activity and dominance behavior in the tube test [30]. Therefore, we compared the firing rate, and other intrinsic neural parameters, with the latency in the tube test (Table S9). We found a significant positive correlation between the spontaneous mPFC spiking activity (firing rate difference, winner – loser) and interaction time (latency difference, winner – loser) in the tube test (P = 0.0142, **Fig. 4D**). This was further confirmed by multiple linear regression (P = 0.0086, Table S9). The correlation was specific to putative pyramidal neurons (P = 0.032, Fig. S13) and was not present in putative interneurons (P = 0.130, Fig. S13). Thus, intrinsic spiking activity in the mPFC correlates with social ranking and dominance behavior. We also explored whether intrinsic cortical activity patterns correlated with animal behavior during spatial navigation. We performed multiple regression analysis including several neural (theta oscillations) and behavioral (task performance) parameters. We did not detect significant regressors among neural factors (Tables S10 and S11). These results show that the activity of mPFC neurons correlates with dominance behavior in the tube test, whereas hippocampal-mPFC connectivity correlates with social behavior in the navigation test. Overall, these data suggest that intrinsic cortical activity and connectivity are significant factors in the regulation of behavioral performance in social contexts.

### Spiking activity in prefrontal cortex correlates with dominance hierarchy and social context

Given the consistent difference in spiking patterns between dominant and subordinate animals under anesthesia, and their relationship with dominance behavior in the social ranking test, we sought to study the spike timing of the mPFC during active behavior. We implanted tetrodes in the mPFC of dominant and submissive mice and recorded their single-unit activity during goal-directed behavior. Here, we further simplified the T-maze protocol, recording only during training sessions and using pairs of animals for collective trials (SI Materials and Methods). The mPFC of freely-moving animals exhibited prominent theta oscillations, similar to those recorded under anesthesia, in both dominant and submissive mice, modulating the spike timing of cortical neurons (**Fig. 5A**). We focused on the firing patterns and theta-phase locking of individual mPFC neurons and compared them across social ranking, behavioral state, and social context (Tables S12 and S13). The firing rates of mPFC neurons were different according to social ranking, with the submissive mice showing larger neuronal activity in the home cage (P = 0.0013, **Fig. 5B**). Similarly, the phase-coupling of mPFC neurons to local theta oscillations was different between social rankings, with the submissive animals exhibiting stronger phase coupling to theta waves during spatial navigation (P = 0.0010, **Fig. 5C**). The social context exerted an effect in both dominant and submissive animals, as the firing rate of mPFC units increased in the presence of the littermate in both cases during navigation (Fig. S14). Given that trial durations were not significantly different between social contexts (P = 0.2217, Fig. S14), this result is unlikely to be explained by different navigation speeds. Similarly, task performance was not different between social contexts, thus this could not be considered a modulating factor of spiking activity (P = 0.2597, Fig. S14). In addition, firing rates of individual neurons were consistent across trials, and exhibited little variation between recording sessions, but maintained a noticeable difference between social contexts (Fig. S14). Next, to directly compare the dynamics of spiking activity during spatial navigation we normalized both firing rates and trial durations. To estimate variability in both dimensions, we computed the variation of both the amplitude and timing of spiking activity (Tables S14 and S15). We found that both parameters were modulated by the significant interaction between dominance hierarchy and social context (hierarchy, P = 0.0102; context, P = 0.0154; Fig. S15). Thus, we plotted and compared the normalized firing rate of mice during spatial navigation. Prefrontal neurons of dominant animals discharged with similar profiles regardless of the social context (P > 0.05, **Fig. 5D**). Conversely, neuronal activity from submissive animals steadily increased during spatial navigation (Fig. S16); particularly during individual trials, which were significantly different from collective trials (P < 0.05, **Fig. 5E**). Overall, these results suggest that task-related mPFC spiking activity correlates with dominance hierarchy and social context.

**Figure 5.**
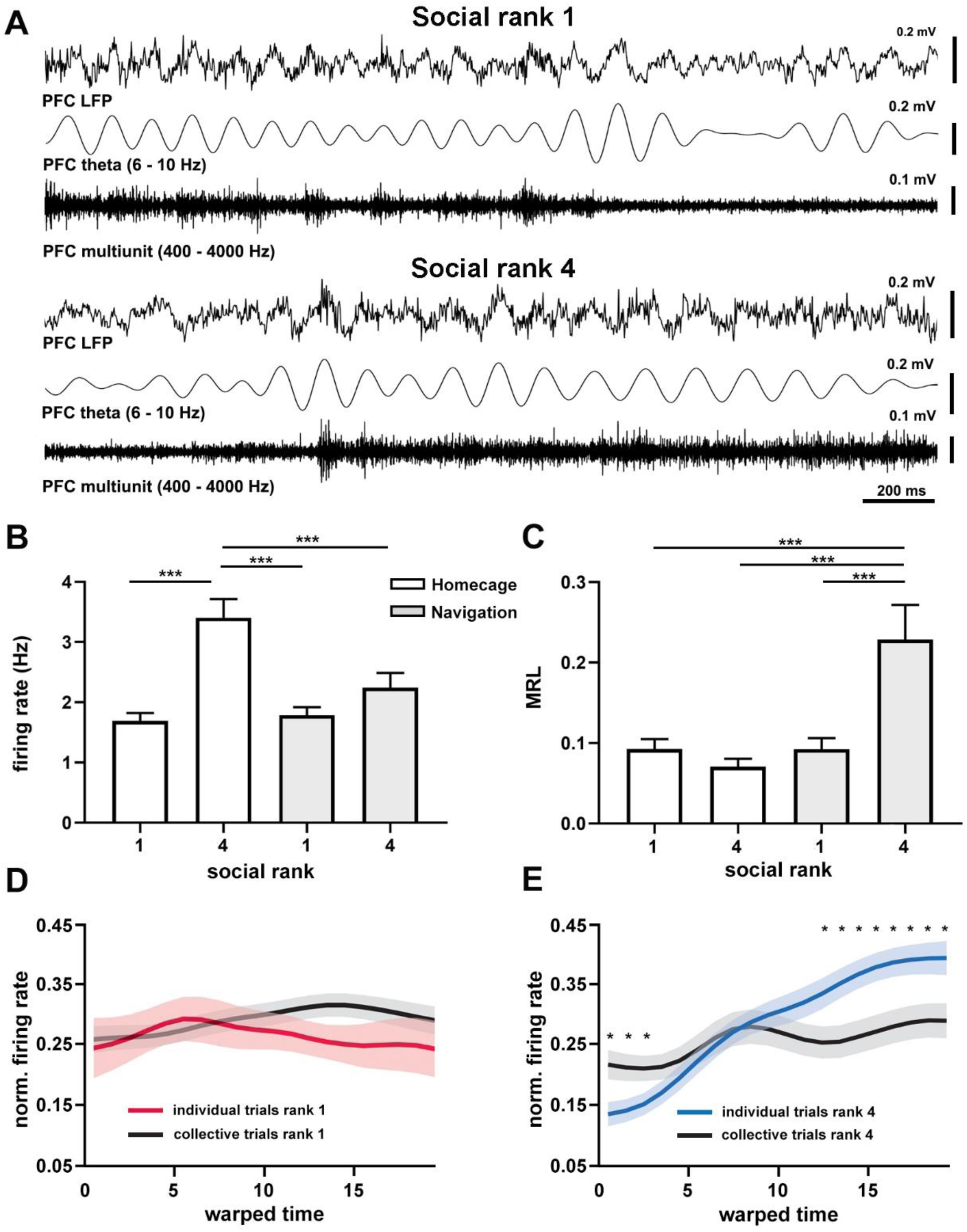
Cortical dynamics of dominance hierarchy during spatial navigation. A, example recordings of prefrontal cortical (LFP PFC) activity showing theta oscillations (filtered 4-8 Hz) and neuronal spiking (units PFC, filtered 300-4000 Hz) from two chronically-implanted dominant (rank 1, mouse H8372) and submissive (rank 4, mouse H8373) mice during spatial navigation in the T-maze. Firing rates (B) and mean resultant length (MRL, C) of mPFC single units (n = 67) according to behavioral state and social ranking. B, two-way ANOVA, P = 0.0013 (state*hierarchy). C, two-way ANOVA, P = 0.0010 (state*hierarchy). Normalized firing rates for dominant (D) and submissive (E) animals during T-maze navigation. Warped time represents entire T-maze. Bin 0, start box; bin 20, arm end. Asterisks depict significant differences between curves, Wilcoxon rank-sum (*, P < 0.05). Bonferroni test post hoc (*, P < 0.05; **, P < 0.01; ***, P < 0.001). Bars, average ± SEM; colored lines, average population; shading areas, ± SEM.

## Discussion

Our results show that goal-directed spatial behavior acquired individually can be disrupted by contingent social interactions during collective navigation. Performance in the social context was critically dependent on both the previous individual learning process, during the training phase, and contingent social interactions arising during navigation, in the testing phase. Ultimately, the influence of contingent social interactions on individual behavior depended on dominance hierarchy and correlated with the intrinsic connectivity of the hippocampal-prefrontal circuit. Moreover, neural spiking in the mPFC partly accounted for social performance during both spatial navigation and dominance behavior. Hence, intrinsic cortical activity and connectivity patterns seemingly differentiate dominance hierarchy and social behavior.

Given that trained animals acquired complete information about the navigation task during the training phase, it might be expected that mice would maintain their stereotyped, efficient navigation strategy during the testing phase of the navigation task. Instead, animals flexibly switched strategies and privileged sensory evidence arising from contingent social interactions, which resulted in significant loss of performance and increased latency during collective movement. The shift in decision-making was not directly related to dominance hierarchy, but to contingent social interactions arising during collective navigation. Indeed, task performance in collective trials was significantly correlated with the distribution of animals in the maze, regardless of the previously learned location of reward during individual trials. Recent studies have shown that as experience increases, mice shift their sensory-based strategy to more efficient, stereotyped foraging based on spatial memory that varies little in response to sensory cues [31]. Conversely, our results show that when sensory experience occurs in social context, it can substantially modify individual behavior. Perceptual evidence arising in the social context is possibly more elaborate than common individual experience as it recruits all sensory modalities and specifically activates circuits for the recognition of and interaction with conspecifics [1]. For example, there is a strongly correlated group structure among mice, as more information about group behavior is contained in the joint position of mice than what can be extracted from summing all the information provided by the interactions between pairs of mice [2, 3]. This irreducible high-order structure in social behavior further supports the study of collective behavior in social groups instead of focusing on individuals.

Ranking systems emerge in social groups to regulate competition over limited resources. Rodents establish dominance hierarchy based on a history of recurrent social interactions in which one subject acts as dominant over the subordinate individuals. Dominance hierarchy can be assessed and quantified by several tests, including the agonistic behavior assay, the barber test, and the ultrasonic test, among others. Here, we used the tube test that is consistent with all those paradigms and has been validated based on transitivity, consistency, and stability [9]. In agreement with previous reports, we found that dominance hierarchy was stable over time [9]. Interestingly, dominance hierarchy had no direct relation with task performance during collective behavior; yet, it distinctly modulated the susceptibility of individuals to contingent social interactions arising during spatial navigation. Previous studies have shown that effective leadership and social decision-making during collective movement do not require intrinsic differences between individuals, such as dominance hierarchy or body size [32]. By studying decision-making during social interactions, we have established that dominant mice do not guide navigation during collective behavior but instead tend to move to the arm that was more densely populated by littermates. This striking behavioral pattern suggests that, in the social context, dominant mice shift their interest from performing the goal-directed task to monitoring the collective behavior of the social group. This pattern is completely at odds with the acquisition of the navigation task, in which dominant and subordinate groups showed comparable performance and latency. Moreover, intrinsic cortical dynamics revealed significant differences according to social ranking, as dominant mice exhibited distinct intrinsic cortical activity and connectivity patterns that segregated them from the subordinate groups, with larger mPFC firing rates, larger hippocampal SWR episodes, weaker coupling between mPFC neurons and cortical theta oscillations, and stronger coupling between mPFC neurons and hippocampal SWR episodes. Intrinsic cortical dynamics, specifically the post-ripple cortical connectivity, correlated with the PSI; that is, the factor of social influence on individual task performance. Overall, cortical connectivity directly correlated with both task performance during individual trials and the social influence on individual behavior. Importantly, these two factors accounted for a large fraction of the variance (62%) in task performance during collective trials.

An important limitation of our study is that we could not track individual trajectories of all littermates during collective trials. Hence, we do not have detailed information about their instantaneous locomotion speed or movement patterns. Cortical oscillations are strongly state-dependent, and lacking such data may be relevant since human and animal experiments support a role for the hippocampus in imagination, planification, and memory retrieval [21, 33]. Importantly, SWRs dominate hippocampal activity during quiescent states [22]; for example, when animals exploring the environment make a pause or stop, and those moments may be highly relevant for temporal prospection or planning [34]. Indeed, hippocampal SWRs precede successful memory retrieval in awake humans [35] and have been proposed to support decision-making and imagination [33]. Our results show that the intrinsic coordinated activity between hippocampal SWRs and prefrontal neuronal spiking was significantly correlated with the PSI. Importantly, hippocampal-prefrontal coordination during SWRs has also been proposed as a neural substrate for decision-making [36]. Indeed, hippocampal spiking during SWRs can represent past or potential future experience [33], and ripple disruptions affect memory performance [34, 37]. Hence, SWRs support both memory consolidation and memory retrieval, which could be at the service of associated cognitive processes such as decision-making. We have previously shown that adverse environmental conditions, such as stress, can impair intrinsic hippocampal-cortical connectivity following SWRs [26]. Importantly, such disruptions are accompanied by alterations in long-term memory [26]. Thus, our current results suggest that the strength of intrinsic hippocampal-cortical connectivity is a potential candidate for the modulation of goal-directed behavior in social groups.

Social relationships can shape individual behavior and affect decision-making [38]. Although dominance hierarchy emerges from recurrent social interactions, it causally results from the synaptic efficacy of excitatory transmission in the mPFC [9]. Hence, dominance hierarchy stems from the matrix of cortical activity and connectivity patterns. The mPFC is essential for decision-making, executive behavior, and social interactions [10]. We report here that intrinsic cortical dynamics expressed in the spontaneous activity of the anesthetized brain partially accounts for dominance hierarchy and social behavior. This suggests that the internally generated, self-organized patterns of cortical activity, unrelated to behavior or relevant perceptual processing, may define the framework of behavioral performance. Naturally, this observation cannot fully account for the shifting in decision-making in the social context, as other cortical regions also contribute to goal-directed spatial navigation. For example, the orbitofrontal cortex is relevant in shifting decisions [39]. Indeed, previous studies have established that orbitofrontal circuits encode the shift between goal-directed and habitual actions [39], thus allowing flexible and efficient decision-making. This is also consistent with recent findings showing that the orbitofrontal cortex integrates prior (i.e., memory) with current (i.e., sensory) signals to guide adaptive behavior [40]. The mPFC exhibits robust anatomical connectivity with the orbitofrontal cortex [10], and these reverberant connections are certainly important in shifting decision-making strategies. Thus, we anticipate that to further understand decision-making in the social context, future studies will have to assess not only ongoing activity of the prefrontal cortex during decision-making, but also the contribution of intrinsic network dynamics in other functionally connected cortical regions.

Finally, to further understand the contribution of individual neurons in the mPFC we recorded their task-related activity during spatial navigation. Interestingly, results from these experiments differed from those obtained under anesthesia, as dominant animals exhibited overall lower and more stable firing rates than submissive animals in the mPFC. This unforeseen outcome could be, at least partially, related with the modified behavioral protocol used in chronically-implanted. Indeed, implanted animals were not food-restricted, did not fully acquire the task, social contexts were tested on different sessions, and most importantly, dominance hierarchy might have changed after surgery (see SI Materials and Methods). Despite these significant variations, we confirmed that spiking patterns of dominant and submissive mice were distinctly modulated by the social context and spatial navigation. Overall, our results suggest that hippocampal-cortical activity and connectivity patterns are important factors to define dominance hierarchy and social behavior. These results suggest that the interplay between contingent social interactions and dominance hierarchy can regulate behavioral performance, supported by the intrinsic matrix of coordinated activity in the hippocampal-prefrontal circuit.

## Supporting information

Tube test rank

Individual acute

Collective acute

Individual chronic

Collective chronic

## Acknowledgments

We thank Daniel Rojas and Fabian Muñoz for commenting on previous versions of the manuscript. This work was funded by Fondecyt regular grant 1190375 (to P.F.); CONICYT, PIA Anillos ACT 172121 (to P.F.); PMD-03/17, Facultad de Medicina, Pontificia Universidad Católica de Chile (to A. L.-V.)

## Materials and methods

Fifteen cages of 4 C57BL/6j littermates (12-25 weeks old) were food-restricted until reaching 85% of ad libitum weight and then trained in a modified T-maze to navigate and find a pellet reward in a fixed pocket. Performance and latency were quantified as the number of correct trials and time interval to reach the reward pocket, respectively. Concomitant with the navigation test, dominance hierarchy was established with the tube test. After behavioral testing, some animals (n = 20) were deeply anesthetized with urethane and the activity of both the hippocampus and prefrontal cortex was monitored. For chronic recordings, 4 cages of 4 C57BL/6j littermates (15 weeks old) were surgically implanted and allowed to recover, after which they were trained in the modified T-maze while recorded in the mPFC. Spectral analysis of LFP coherence and power were computed using multitaper Fourier analysis from the Chronux toolbox (http://www.chronux.org) by using MatLab. Spike sorting was performed offline using MATLAB based graphical cluster-cutting software, Mclust/Klustakwik-toolbox (version 3.5, [45]) and phase-locking analysis was computed using the Matlab toolbox CircStats (http://philippberens.wordpress.com/code/circstats/). To verify recording sites, Nissl-staining was conducted following the completion of study. See SI Appendix for a detailed description of experimental and analytic methods.

## Notes

### Competing Interest Statement

The authors have declared no competing interest.

## References

1. Insel TR, Fernald RD (2004) How the brain processes social information: searching for the social brain. Annu Rev Neurosci 27:697–722. https://doi.org/10.1146/annurev.neuro.27.070203.144148

2. Shemesh Y, Sztainberg Y, Forkosh O, et al (2013) High-order social interactions in groups of mice. Elife 2:e00759. https://doi.org/10.7554/eLife.00759

3. de Chaumont F, Coura RD-S, Serreau P, et al (2012) Computerized video analysis of social interactions in mice. Nat Methods 9:410–417. https://doi.org/10.1038/nmeth.1924

4. Conradt L, Roper TJ (2003) Group decision-making in animals. Nature 421:155–158. https://doi.org/10.1038/nature01294

5. Lindzey G, Winston H, Manosevitz M (1961) Social Dominance in Inbred Mouse Strains. Nature 191:474. https://doi.org/10.1038/191474a0

6. Lathe R (2004) The individuality of mice. Genes Brain Behav 3:317–327. https://doi.org/10.1111/j.1601-183X.2004.00083.x

7. Park M-J, Seo BA, Lee B, et al (2018) Stress-induced changes in social dominance are scaled by AMPA-type glutamate receptor phosphorylation in the medial prefrontal cortex. Sci Rep 8:15008. https://doi.org/10.1038/s41598-018-33410-1

8. Anacker AMJ, Moran JT, Santarelli S, et al &Enhanced Social Dominance and Altered Neuronal Excitability in the Prefrontal Cortex of Male KCC2b Mutant Mice. Autism Research 0:https://doi.org/10.1002/aur.2098

9. Wang F, Zhu J, Zhu H, et al (2011) Bidirectional control of social hierarchy by synaptic efficacy in medial prefrontal cortex. Science 334:693–697. https://doi.org/10.1126/science.1209951

10. Euston DR, Gruber AJ, McNaughton BL (2012) The role of medial prefrontal cortex in memory and decision making. Neuron 76:1057–1070. https://doi.org/10.1016/j.neuron.2012.12.002

11. Matsumoto M, Matsumoto K, Abe H, Tanaka K (2007) Medial prefrontal cell activity signaling prediction errors of action values. Nat Neurosci 10:647–656. https://doi.org/10.1038/nn1890

12. Benchenane K, Peyrache A, Khamassi M, et al (2010) Coherent theta oscillations and reorganization of spike timing in the hippocampal-prefrontal network upon learning. Neuron 66:921–936. https://doi.org/10.1016/j.neuron.2010.05.013

13. Lee E, Rhim I, Lee JW, et al (2016) Enhanced Neuronal Activity in the Medial Prefrontal Cortex during Social Approach Behavior. J Neurosci 36:6926–6936. https://doi.org/10.1523/JNEUROSCI.0307-16.2016

14. Olton DS (1979) Mazes, maps, and memory. Am Psychol 34:583–596

15. Couzin ID, Krause J, James R, et al (2002) Collective memory and spatial sorting in animal groups. J Theor Biol 218:1–11

16. Partridge BL, Pitcher T, Cullen JM, Wilson J (1980) The three-dimensional structure of fish schools. Behav Ecol Sociobiol 6:277–288. https://doi.org/10.1007/BF00292770

17. Partridge BL (1982) The structure and function of fish schools. Scientific American 246:114–123. https://doi.org/10.1038/scientificamerican0682-114

18. Wang F, Kessels HW, Hu H (2014) The mouse that roared: neural mechanisms of social hierarchy. Trends Neurosci 37:674–682. https://doi.org/10.1016/j.tins.2014.07.005

19. Negrón-Oyarzo I, Espinosa N, Aguilar-Rivera M, et al (2018) Coordinated prefrontal-hippocampal activity and navigation strategy-related prefrontal firing during spatial memory formation. Proc Natl Acad Sci USA 115:7123–7128. https://doi.org/10.1073/pnas.1720117115

20. Buzsáki G, Moser EI (2013) Memory, navigation and theta rhythm in the hippocampal-entorhinal system. Nat Neurosci 16:130–138. https://doi.org/10.1038/nn.3304

21. Buzsáki G (2015) Hippocampal sharp wave-ripple: A cognitive biomarker for episodic memory and planning. Hippocampus 25:1073–1188. https://doi.org/10.1002/hipo.22488

22. Ylinen A, Bragin A, Nádasdy Z, et al (1995) Sharp wave-associated high-frequency oscillation (200 Hz) in the intact hippocampus: network and intracellular mechanisms. J Neurosci 15:30–46

23. Remondes M, Wilson MA (2015) Slow-γ Rhythms Coordinate Cingulate Cortical Responses to Hippocampal Sharp-Wave Ripples during Wakefulness. Cell Rep 13:1327–1335. https://doi.org/10.1016/j.celrep.2015.10.005

24. Logothetis NK, Eschenko O, Murayama Y, et al (2012) Hippocampal-cortical interaction during periods of subcortical silence. Nature 491:547–553. https://doi.org/10.1038/nature11618

25. Siapas AG, Wilson MA (1998) Coordinated interactions between hippocampal ripples and cortical spindles during slow-wave sleep. Neuron 21:1123–1128

26. Negrón-Oyarzo I, Neira D, Espinosa N, et al (2015) Prenatal Stress Produces Persistence of Remote Memory and Disrupts Functional Connectivity in the Hippocampal-Prefrontal Cortex Axis. Cereb Cortex 25:3132–3143. https://doi.org/10.1093/cercor/bhu108

27. Buzsáki G (2002) Theta oscillations in the hippocampus. Neuron 33:325–340

28. Jones MW, Wilson MA (2005) Theta Rhythms Coordinate Hippocampal–Prefrontal Interactions in a Spatial Memory Task. PLoS Biol 3:. https://doi.org/10.1371/journal.pbio.0030402

29. Hagan MA, Dean HL, Pesaran B (2012) Spike-field activity in parietal area LIP during coordinated reach and saccade movements. J Neurophysiol 107:1275–1290. https://doi.org/10.1152/jn.00867.2011

30. Zhou T, Zhu H, Fan Z, et al (2017) History of winning remodels thalamo-PFC circuit to reinforce social dominance. Science 357:162–168. https://doi.org/10.1126/science.aak9726

31. Gire DH, Kapoor V, Arrighi-Allisan A, et al (2016) Mice Develop Efficient Strategies for Foraging and Navigation Using Complex Natural Stimuli. Curr Biol 26:1261–1273. https://doi.org/10.1016/j.cub.2016.03.040

32. Couzin ID, Krause J, Franks NR, Levin SA (2005) Effective leadership and decision-making in animal groups on the move. Nature 433:513–516. https://doi.org/10.1038/nature03236

33. Joo HR, Frank LM (2018) The hippocampal sharp wave-ripple in memory retrieval for immediate use and consolidation. Nat Rev Neurosci 19:744–757. https://doi.org/10.1038/s41583-018-0077-1

34. Jadhav SP, Kemere C, German PW, Frank LM (2012) Awake hippocampal sharp-wave ripples support spatial memory. Science 336:1454–1458. https://doi.org/10.1126/science.1217230

35. Vaz AP, Inati SK, Brunel N, Zaghloul KA (2019) Coupled ripple oscillations between the medial temporal lobe and neocortex retrieve human memory. Science 363:975–978. https://doi.org/10.1126/science.aau8956

36. Yu JY, Frank LM (2015) Hippocampal-cortical interaction in decision making. Neurobiol Learn Mem 117:34–41. https://doi.org/10.1016/j.nlm.2014.02.002

37. Girardeau G, Benchenane K, Wiener SI, et al (2009) Selective suppression of hippocampal ripples impairs spatial memory. Nat Neurosci 12:1222–1223. https://doi.org/10.1038/nn.2384

38. Torquet N, Marti F, Campart C, et al (2018) Social interactions impact on the dopaminergic system and drive individuality. Nat Commun 9:3081. https://doi.org/10.1038/s41467-018-05526-5

39. Gremel CM, Costa RM (2013) Orbitofrontal and striatal circuits dynamically encode the shift between goal-directed and habitual actions. Nat Commun 4:2264. https://doi.org/10.1038/ncomms3264

40. Nogueira R, Abolafia JM, Drugowitsch J, et al (2017) Lateral orbitofrontal cortex anticipates choices and integrates prior with current information. Nat Commun 8:14823. https://doi.org/10.1038/ncomms14823

41. Aasebø IEJ, Lepperød ME, Stavrinou M, et al (2017) Temporal Processing in the Visual Cortex of the Awake and Anesthetized Rat. eNeuro 4:. https://doi.org/10.1523/ENEURO.0059-17.2017

42. Steriade M, Timofeev I, Grenier F (2001) Natural waking and sleep states: a view from inside neocortical neurons. J Neurophysiol 85:1969–1985. https://doi.org/10.1152/jn.2001.85.5.1969

43. Noda T, Takahashi H (2015) Anesthetic effects of isoflurane on the tonotopic map and neuronal population activity in the rat auditory cortex. European Journal of Neuroscience 42:2298–2311. https://doi.org/10.1111/ejn.13007

44. Chauvette S, Crochet S, Volgushev M, Timofeev I (2011) Properties of Slow Oscillation during Slow-Wave Sleep and Anesthesia in Cats. J Neurosci 31:14998–15008. https://doi.org/10.1523/JNEUROSCI.2339-11.2011

45. Harris KD, Henze DA, Csicsvari J, et al (2000) Accuracy of tetrode spike separation as determined by simultaneous intracellular and extracellular measurements. J Neurophysiol 84:401–414. https://doi.org/10.1152/jn.2000.84.1.401

46. Mitra PP, Pesaran B (1999) Analysis of dynamic brain imaging data. Biophys J 76:691–708. https://doi.org/10.1016/S0006-3495(99)77236-X

