## Supplemental information for "Cortical dynamics underlying social behavior in dominance hierarchy and spatial navigation"

**This PDF file contains:**

SI Appendix

- SI Methods
- SI References
- SI Figures 1 to 16
- SI Tables 1 to 16

**Other supplementary materials for this manuscript include the following:**

Movies S1 to S5

**SI Appendix**

**1. SI Methods**

**Ethics statement**

Efforts were performed to minimize the number of animals used and their suffering. Experimental procedures related to animals were approved by the Institutional Animal Ethics Committee of the Pontificia Universidad Catolica de Chile (protocol code CEBA-13-040).

**Animals**

Groups of four male sibling mice (C57BL/6J strain, n = 19 cages, 20–30 g) were reared together after weanling. All tests were conducted between 10.00 a.m. and 4.00 p.m. Animals were housed under controlled temperature (22 ± 1°C) and humidity (50%) conditions with food and water ad libitum. A 12 h:12 h light–dark cycle was maintained throughout experiments, lights being on from 8.00 a.m. to 8.00 p.m.

**Tube test**

We measured hierarchical relations within social groups with the tube test [1]. Each mouse was placed at the ends of a narrow tube facing inward and one mouse forces the other to back out of the tube with score 1 for the winner and 0 for the loser per session [2]. All mice were tested pairwise for dominance for ten consecutive days using a round-robin design, and the social rank was assessed based on winning against the other cage mates.

**T-maze test**

We used transparent Plexiglas to manufacture the T-maze. For habituation, animals (15 cages of 4 littermates) were individually placed in the maze and allowed to explore freely for 3-5 min for two days. Crumbs of sugared cereal were randomly distributed throughout the maze to stimulate exploration. Animals were then food-restricted to enhance exploration and learning of the arm baited with food, with 1.5 gr of pellet per animal per day. This treatment produced a general weight loss of about 20% in most animals. During the individual phase, we trained animals individually to look for food in a fixed location, at the end of the maze arms. Each animal had 10 trials per day, with a maximum duration of 90 second per trial. During the first trials, if the mouse did not explore, it was gently pushed towards the baited arm. At the end of every trial, the mouse was placed back in the homecage with its siblings and the maze was quickly wiped with 10% EtOH to remove odour cues. In this way, the inter-trial interval for every mouse was around 10 minutes. For every trial we computed the latency or the time interval that every animal took to get from the start box to reach the food reward in the baited arm. In addition, we calculated the performance as the proportion of correctly performed trials based on the first decision to turn to the baited arm. Training finalized when animals reached the learning criterion, meaning that at least three out of four littermates in the box performed correctly six out of eight trials (75%) on two consecutive days. After reaching the learning criterion, we started the collective phase. Every animal performed four individual trials and the next trial was collective, with all four littermates placed in the start box. This was repeated twice, so to complete 10 total trials, 8 individual and 2 collective per day, during 5 consecutive days. During collective trials, reward was randomly assigned according to four options: right arm, left arm, both arms, or none. Videos of individual and collective tests were scored manually on a frame-by-frame basis. The experimenter was blind to the animal identity during scoring.

For chronically-implanted animals (4 cages of 4 littermates), we modified the behavioral protocol due to the cranial implants. Each littermate was trained for 5-8 days individually, while performing the tube test to establish dominance hierarchy. Animals were food-deprived 3-4 hours before each test and this affected performance as mice rarely reached the learning criterium. After the surgery, performed only on dominant and submissive animals, each mouse was isolated from the other littermates for recovery and to prevent removal of the cranial implant. After recovery, animals were re-trained for 3-4 days individually. Then, for electrophysiological recordings, the training and testing phases were separated in two consecutive days. Indeed, eight individual trials were alternately tested in each animal, and the day after eight consecutive collective trials were performed. During these individual and collective trials in the T-maze, animals were connected to the amplifier and cortical neural activity was recorded. Furthermore, before experiments animals were housed together in the same homecage in the morning to habituate them to the presence of the littermate, and tests were performed in the afternoon, after 4-6 hours. Dominance hierarchy could not be established after the surgery due to the cranial implants.

**Acute surgery**

After finishing collective testing, animals were allowed to recover ad libitum weight and then were used for electrophysiological recordings. We recorded simultaneous neuronal activity in the prefrontal cortex and hippocampus. Animals were anesthetized with urethane (0.8 g/kg dissolved in saline, i.p.) and a mixture of ketamine/xylazine (40 mg/kg ketamine; 4 mg/kg xylazine dissolved in saline, i.p.). Anaesthesia was maintained throughout the experiment with urethane administered every 20 minutes with a bomb when required. During the entire experiment, glucosamine solution (0.5-1 ml) was injected subcutaneously every 2 hours to maintain the animal hydrated and body temperature was maintained at 36 ± 1°C using a homeothermic blanket (Harvard Apparatus, MA, USA) and monitored with a rectal probe connected to a temperature controller (Harvard Apparatus, MA, USA). Animals were firmly placed in a stereotaxic frame (Stoelting Co.).

**Acute recordings**

To simultaneously record neuronal activity of the prefrontal cortex (cingulate and prelimbic cortex) and the CA1 area of the dorsal hippocampus, small craniotomies (1 mm) were drilled on the skull (right hemisphere) over the recording sites. The stereotaxic coordinates, indicated by the stereotaxic atlas (Franklin and Paxinos 2007), were (relative to bregma): prefrontal cortex, anteroposterior, +2 mm; mediolateral, +0.5 mm; and CA1 hippocampus, anteroposterior, −3 mm; mediolateral, +1.7 mm. The electrodes were slowly lowered via a motorized microdrive (Siskiyou, Grants Pass, OR, USA) to the recording positions. The electrodes were positioned at ∼1.0–2.0 mm dorsoventrally to record activity in the PFC and to record in the CA1 the electrodes were placed at ∼1.1 mm dorsoventrally using the firing of CA1 pyramidal cells and the appearance of SWR as the hallmark for functional localization of the hippocampus. Neuronal activity in the prefrontal cortex was recorded extracellularly with a 32 channel-two shank silicon probe (Poly A32, Neuronexus, mean site resistance ~1 MΩ) stained with DiI. Neuronal activity in the hippocampus was recorded with a 16 channel-silicon probe (A16, Neuronexus, mean site resistance ~1 MΩ) stained with DiI and inserted into the brain with a 30° angle through the midline. Electrical activity was acquired with a 32-channel Intan RHD 2132 amplifier board connected to an RHD2000 evaluation system (Intan Technologies). Single-unit activity and local field potential (LFP; sampling rate 20 kHz) were digitally filtered between 300 Hz –5 kHz and 0.3 Hz – 2 kHz, respectively. Once a spiking multiunit was detected, the simultaneous prefrontal cortex and hippocampal recording started, and lasted for 10 min.

**Chronic surgery**

A survival surgery was conducted on adult mice (3–6 months of age), weighing 21–25 g. Animals were anesthetized with 1.5–2.5% isoflurane, delivered with O2 (2 L per min), and were placed in a stereotaxic frame. Using a surgical drill (Foredom Electric, Bethel, CT), two craniotomies were made, one on the occipital bone for a brass ground screw and another above the frontal bone for the tetrode array. The dura mater was removed for the following craniotomy. Mice were implanted with customized microdrives weighing ~1.5 g, including the dental cement. The array was targeted to the right PFC area, using the coordinates anteroposterior, +2 mm; mediolateral, +0.5 mm, and lowered to an initial depth of 1500 µm below the brain surface. The exposed craniotomy was filled with sterile Vaseline. As the dental acrylic hardened, mice were injected with buprenorphine (0.05 mg/kg) subcutaneously prior to the removal of anesthesia. Mice were closely monitored for the first few hours post-surgery and were monitored daily thereafter

**Chronic recordings**

We recorded neural activity via a unitary gain headstage preamplifier (HS-18; Neuralynx Bozeman, MT), which was connected to an amplifier (Cheetah 32, Neuralynx) linked to the acquisition software (Cheetah 32, Neuralynx). Single units were recorded at a sampling rate of 30 kHz, band-pass filtered (600–6 kHz), and referenced to a nearby 50-µm local reference electrode in corpus callosum above dorsal CA1. Local field potentials were also acquired at a sampling rate of 3 kHz, band-pass filtered (0.1–6 kHz), and referenced to a ground screw above the cerebellum. This was accomplished by a video camera, mounted above the chamber, to the video input of the Cheetah software.

**Histology**

At the end of the electrophysiological recording mice were immediately perfused with 20 ml of saline solution followed by 50 ml of 4% paraformaldehyde in phosphate buffered saline (PBS, pH = 7.4). The brain was removed, incubated overnight in 4% paraformaldehyde in PBS buffer and then stored in PBS buffer containing 0.2% sodium azide. Coronal brain slices (60 -80 µm) were prepared from paraformaldehyde-fixed brains with a vibratome (World Precision Instruments, Sarasota, USA) in ice-cold PBS buffer. For visualization, slices were washed three times in PBS buffer at room temperature and then placed on slides using a mounting medium (Dako) and then, were stained with Nissl-staining, Images were acquired with an epifluorescence microscope for DiI labelling and Nissl-staining (Nikon eclipse Ci).

**Spike sorting**

Neuronal spikes were extracted from prefrontal cortex recordings using semiautomatic clustering KlustaKwik (https://github.com/kwikteam/klustakwik2/). This method was applied over the 32 channels of the silicon probe, grouped in eight pseudo-tetrodes of four nearby channels. Spike clusters were considered single units if their auto-correlograms had a 2-ms refractory period, and their cross-correlograms with all other clusters did not have sharp peaks within 2 ms of 0 lag.

**T-maze trial-warping**

Individual and collective trials were linearly time-warped to produce equal numbers of data points between the start box and the baited arm. In each trial, the number of bins was set to 20, and the firing rate was normalized to its peak value, with a threshold of at least 10 spikes. Finally, data was smoothed with a Gaussian-weighted moving average filter (smoothdata function in MATLAB).

**Brain-state and time-frequency analysis**

We defined brain-states based on the hippocampal LFP. Decomposition of LFP in PFC and hippocampus was performed with multi-taper Fourier analysis [3] implemented in Chronux toolbox (http://www.chronux.org). LFP was downsampled to 500 Hz before decomposition. We recognized theta oscillations, non-theta epochs, and ripple episodes. Unless stated, the LFP from dorsal CA1 stratum pyramidal was considered as the time-frame reference for the spike-timing of recorded cells.

Theta oscillations were detected by calculating the continuous ratio between the envelopes of theta (4–8 Hz) and delta (2–3 Hz) frequency bands filtered from the hippocampus LFP and calculated by the Hilbert transform. A ratio of 1.4 SD or higher, during at least 2 s defined epochs of theta oscillations. Recording episodes outside theta oscillations were defined as non-theta epochs.

Sharp wave-ripples were recorded in dorsal CA1, as close as possible to stratum pyramidale and considered as the time-frame reference for the spike-timing of the recorded neurons and population activity (LFP) in prefrontal cortex. We used a recently described method for ripples detection [4] with some modifications. Briefly, the hippocampus LFP was first down-sampled to 1 kHz, then band-pass filtered (100-200 Hz) using a zero-phase shift non-causal finite impulse filter with 0.5 Hz roll-off. Next, the signal was rectified, and low-pass filtered at 20 Hz with a 4th order Butterworth filter. This procedure yields a smoothed envelope of the filtered signal, which was then z-score normalized using the mean and SD of the whole signal in the time domain. Epochs during which the normalized signal exceeds a 3.5 SD threshold were considered as ripple events. The first point before threshold that reached 1 SD was considered the onset and the first one after threshold to achieve 1 SD as the end of events. The difference between onset and end of events was used to estimate the ripple duration. We introduced a 50 ms-refractory window to prevent double detections. To precisely determine the mean frequency, amplitude, and duration of each event, we performed a spectral analysis using Morlet complex wavelets of seven cycles. The Matlab toolbox used is available online as LAN toolbox (http://lantoolbox.wikispaces.com/).

**Cross-correlation analysis**

Activity of spiking neurons and hippocampal ripples was cross-correlated by applying the "sliding-sweeps" algorithm [5]. A time window of ± 1 s was defined with the 0-point assigned to the start time of a ripple. The timestamps of the cortical spikes within the time window were considered as a template and were represented by a vector of spikes relative to t = 0 s, with a time bin of 50 ms and normalized to the total number of spikes. Thus, the central bin of the vector contained the ratio between the number of PFC spikes elicited between ± 25 ms and the total number of spikes within the template. Next, the window was shifted to successive ripples throughout the recording session, and an array of recurrences of templates was obtained. Both prefrontal cortex timestamps and start times of ripples where shuffled (1000 samples) samples by randomized exchange of the original inter-event intervals and the cross-correlation procedure was performed on the pseudo-random sequence.

**Theta phase locking**

Phase-locking analysis was computed using the Matlab toolbox CircStats (http://philippberens.wordpress.com/code/circstats/). Briefly, LFP traces were bandpass filtered in the theta range (4-8 Hz in anesthesia and 6-10 Hz in freely-moving animals, respectively; with zero phases shift non-causal finite impulse filter with 0.5 Hz roll-off). Phase locking was quantified as the circular concentration of the resulting phase distribution, which was defined as resultant mean length ($\mathrm{MRL} = \sqrt{\mathrm{Z/n}}$) where Z is calculated from Rayleigh test and n is the number of spikes [6]. The statistical significance of phase-locking was assessed using the Rayleigh test for circular uniformity. To avoid bias, we only considered neurons with >20 action potentials.

**Statistics**

We performed inter-subject comparisons to establish if behaviour and simultaneous cortico-hippocampal activity were different across social rank. We pooled neuronal data from all animal of a specific social rank in the same experimental group for all other statistical analysis. Data were tested for normality using the Kolmogorov–Smirnov test and then compared with the appropriate test with parametric analysis (one-way ANOVA followed by Bonferroni post-hoc test). Comparison between behavioural parameters and other non-normally distributed parameters were analysed with non-parametric tests (Wilcoxon signed rank; Kruskal-Wallis test followed by Dunn's multiple comparisons post-hoc test). The statistical significance of the observed repetition of spike sequences was assessed by comparing, bin to bin, the original sequence with the shuffled sequence. An original correlation sequence that presented a statistical distribution different from 1000 permutations was considered as statistically significant, with $P < 0.01$ probability, instead of a chance occurrence. Linear correlations between parameters were analysed by Spearman correlation test. To calculate the P-value we used the circ_corrcl.m in the CircStat toolbox of MATLAB (The Mathworks, Inc.) and STATISTICA 7.0 software (StatSoft, Inc). Summary results of statistical tests are presented in Table S16.

### 2. SI References

1. Lindzey G, Winston H, Manosevitz M (1961) Social Dominance in Inbred Mouse Strains. Nature 191:474.
2. Wang F, Zhu J, Zhu H, et al. (2011) Bidirectional control of social hierarchy by synaptic efficacy in the medial prefrontal cortex. Science 334:693–697.
3. Mitra, P. P. & Pesaran, B. Analysis of dynamic brain imaging data. *Biophys J* **76**(2), 691–708 (1999).
4. Logothetis NK, Eschenko O, Murayama Y, et al. (2012) Hippocampal-cortical interaction during periods of subcortical silence. Nature 491:547–553.
5. Abeles M & Gerstein GL (1988) Detecting spatiotemporal firing patterns among simultaneously recorded single neurons. Journal of Neurophysiology 60(3):909—924.
6. Adhikari A, Topiwala MA, & Gordon JA (2011) Single units in the medial prefrontal cortex with anxiety-related firing patterns are preferentially influenced by ventral hippocampal activity. Neuron 71(5):898--910.

### 3. SI Figures

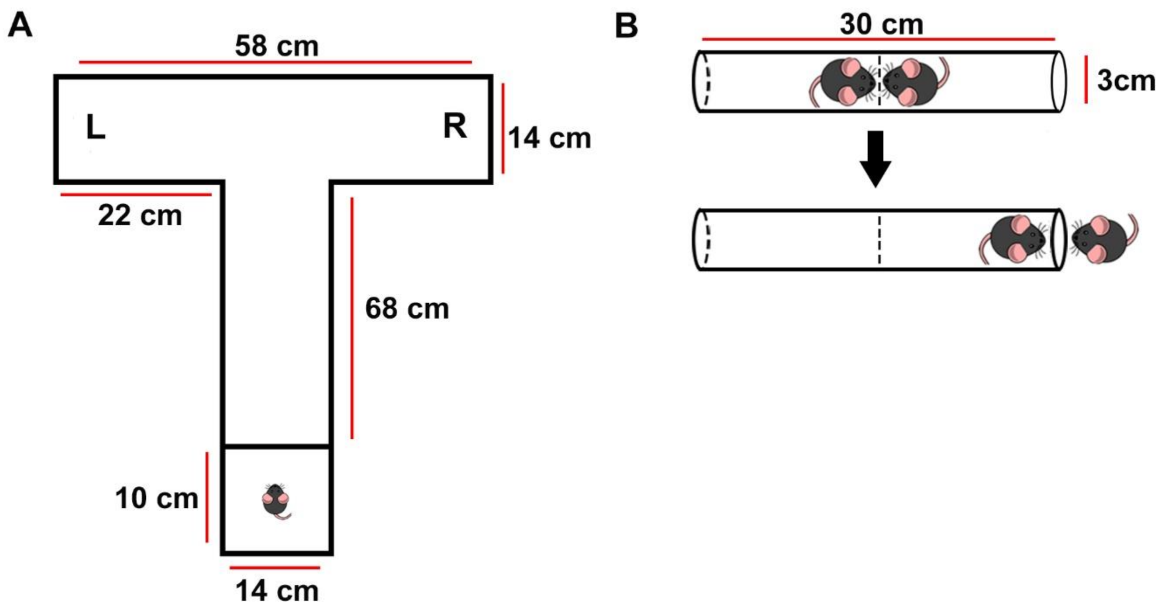

**Figure S1.** Behavioral apparatuses (A) T-maze (B) tube test.

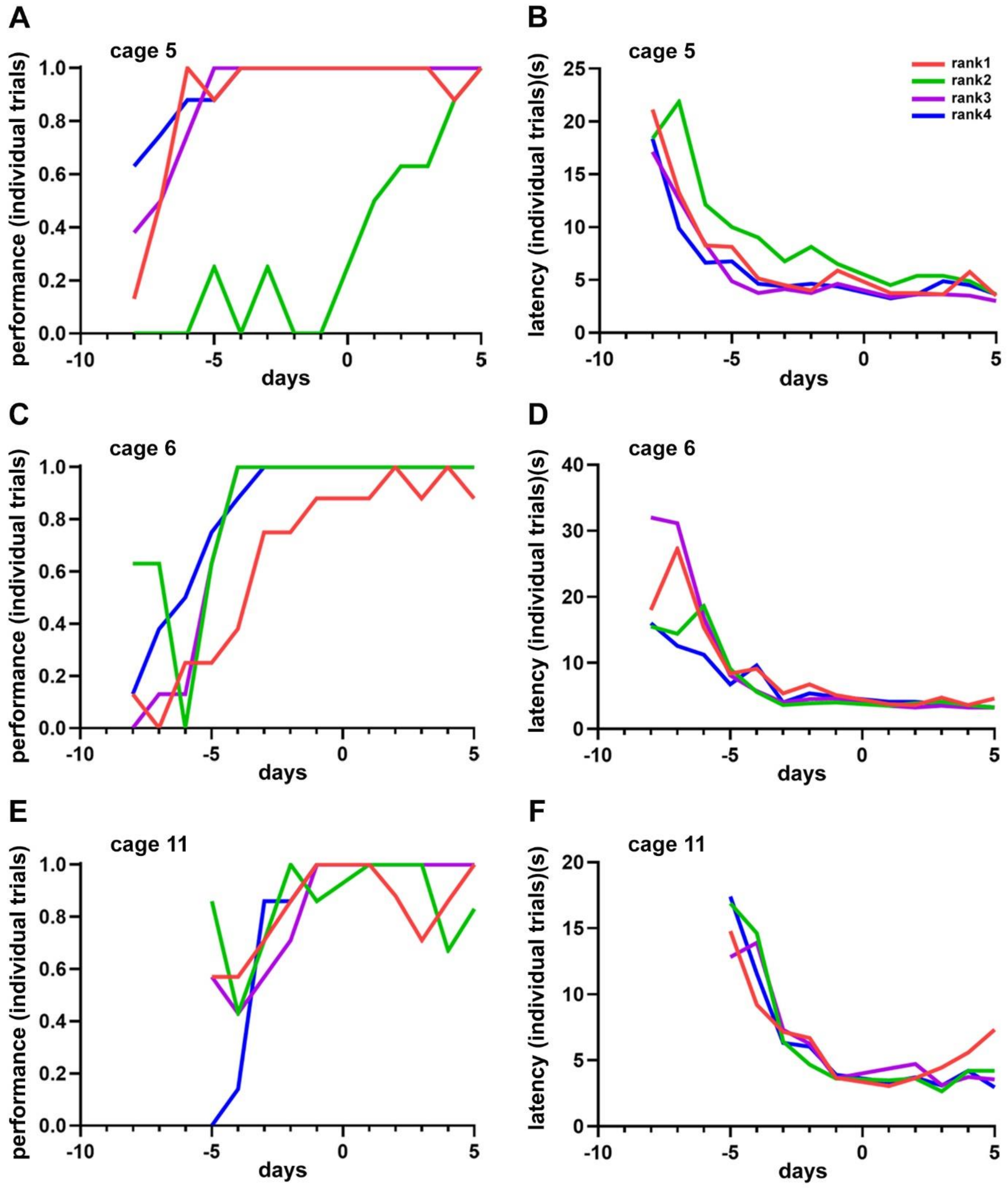

**Figure S2.** Example learning curves showing performance (A, C, E) and latency (B, D, F) for cages of littermates sorted by social ranking (color coded). Colored lines, average trials.

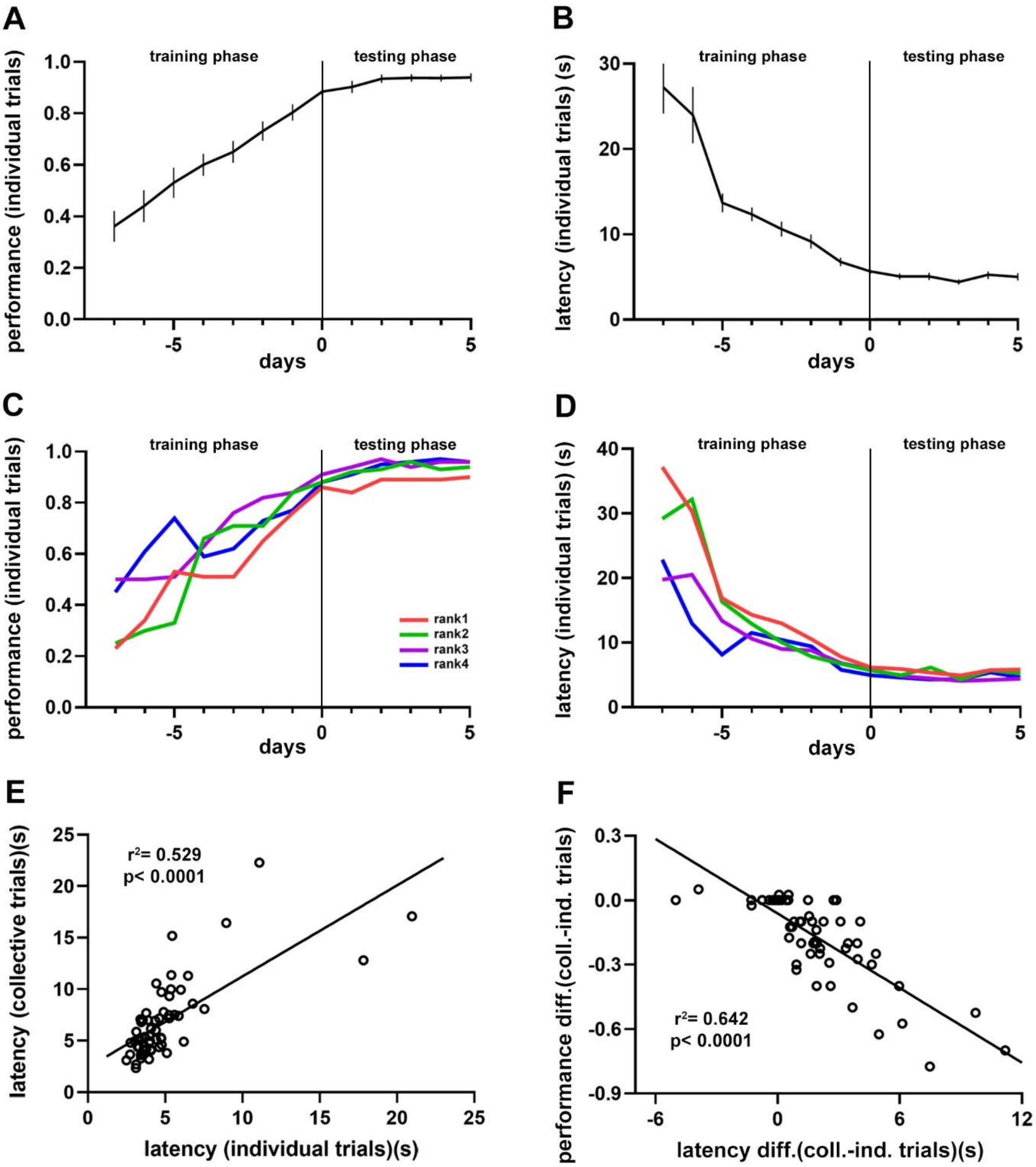

**Figure S3.** Average performance (A) and latency (B) during training and testing phases for all tested animals (n = 60). Data are presented as mean ± SEM. Average performance (C) and latency (D) during both training and testing phases for all tested animals sorted by social ranking (color coded). E, average latency from individual trials against average latency from collective trials. F, average latency difference (collective – individual trials) against average performance difference (collective – individual trials). Black lines, population averages ± SEM; colored lines, average population.

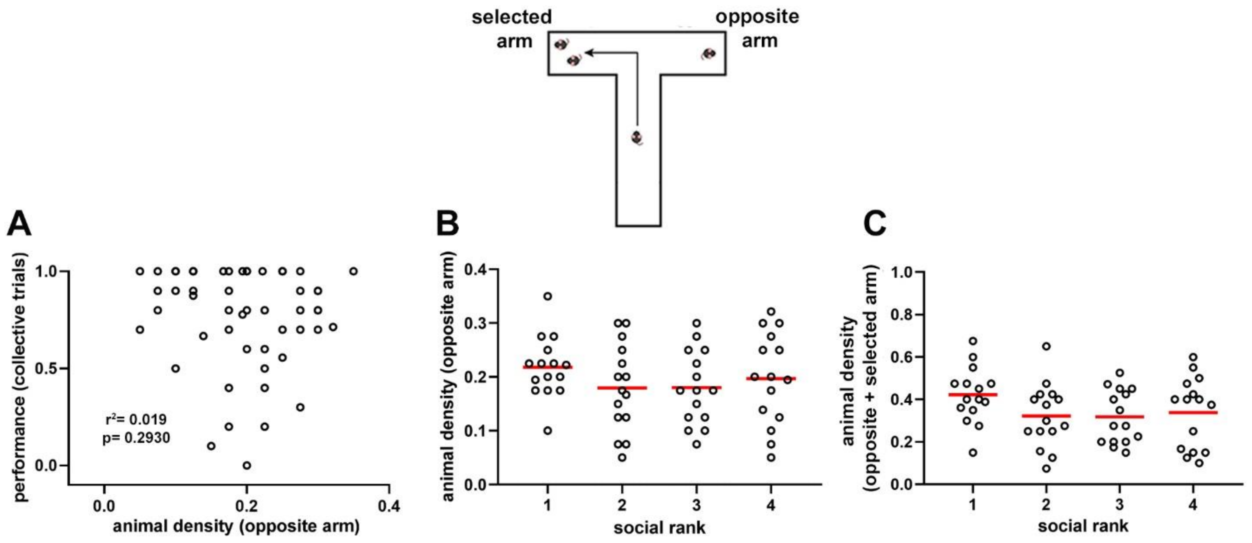

**Figure S4.** Density of animals in the lateral arms during collective navigation. A, Scatter plot of average animal density in the opposite arm and performance of individual mice during collective trials (n = 60). Linear regression was not statistically significant. B, density of animals located in the opposite arm to the animal's decision according to social ranking. One-way ANOVA, P = 0.4499. C, average density of animals located in both lateral arms when the choosing mouse was located at the junction according to social ranking. One-way ANOVA, P = 0.1783. Circles, individual mice average; red line; population average.

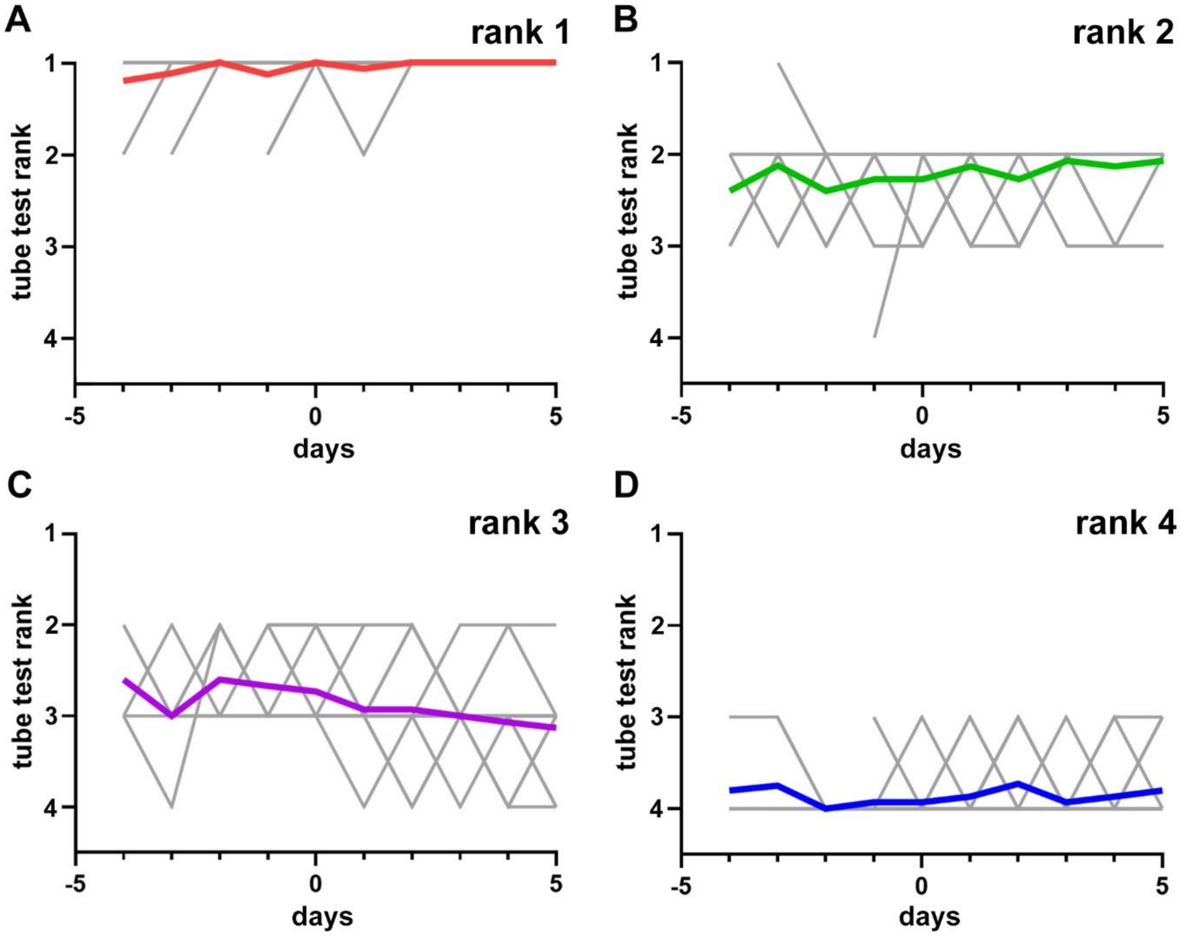

**Figure S5.** Dominance hierarchy of animals performing the spatial navigation task. Summary plots for all measured animals (n = 60) according to social ranking. Days -5 to 0, training phase; days 1 to 5, testing phase. A, ranking 1, dominant; B, ranking 2, first active subordinate; C, ranking 3, second active subordinate; D, ranking 4, submissive. Colored lines, population average; gray lines, individual mice. Note ranking stability over time, particularly for dominant mice.

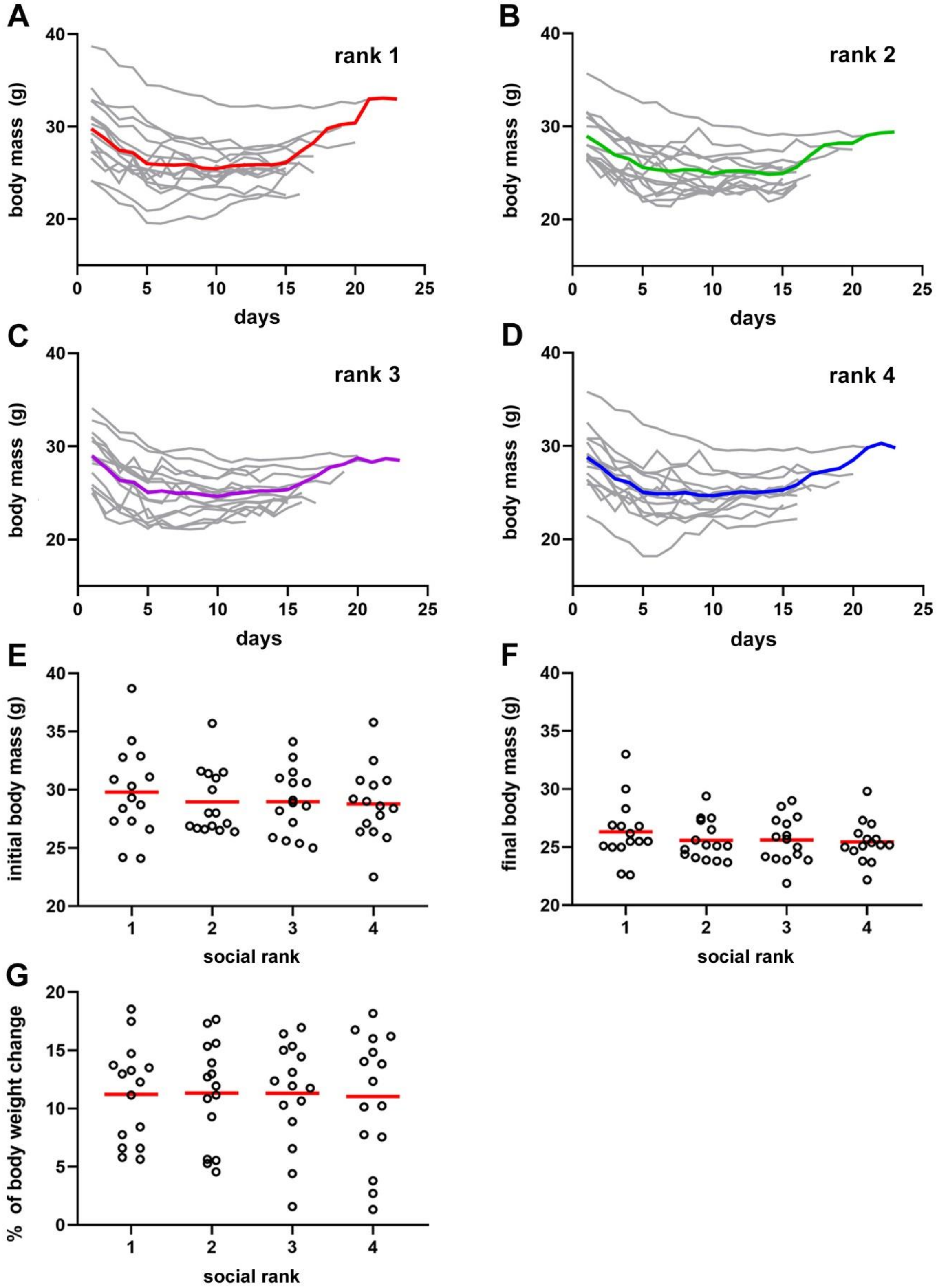

**Figure S6.** Body weight of animals performing the spatial navigation task. Day 0 represents initial weight. A, ranking 1, dominant; B, ranking 2, first active subordinate; C, ranking 3, second active subordinate; D, ranking 4, submissive. Food was restricted from day 1 for every mouse until it reached roughly 85% of its original weight. Summary plots for all tested animals (n = 60) according to social ranking with free access to food (E) and during food restriction (F). Note that dominant animals do not exhibit different body mass when compared to the other social rankings. G, maximal weight loss during food restriction protocol. E, One-way ANOVA, P = 0.8216; F, One-way ANOVA, P

= 0.6554. G, One-way ANOVA, P = 0.983. Colored lines, average population; gray lines, individual mice; circles, average of individual animals; red lines; population average.

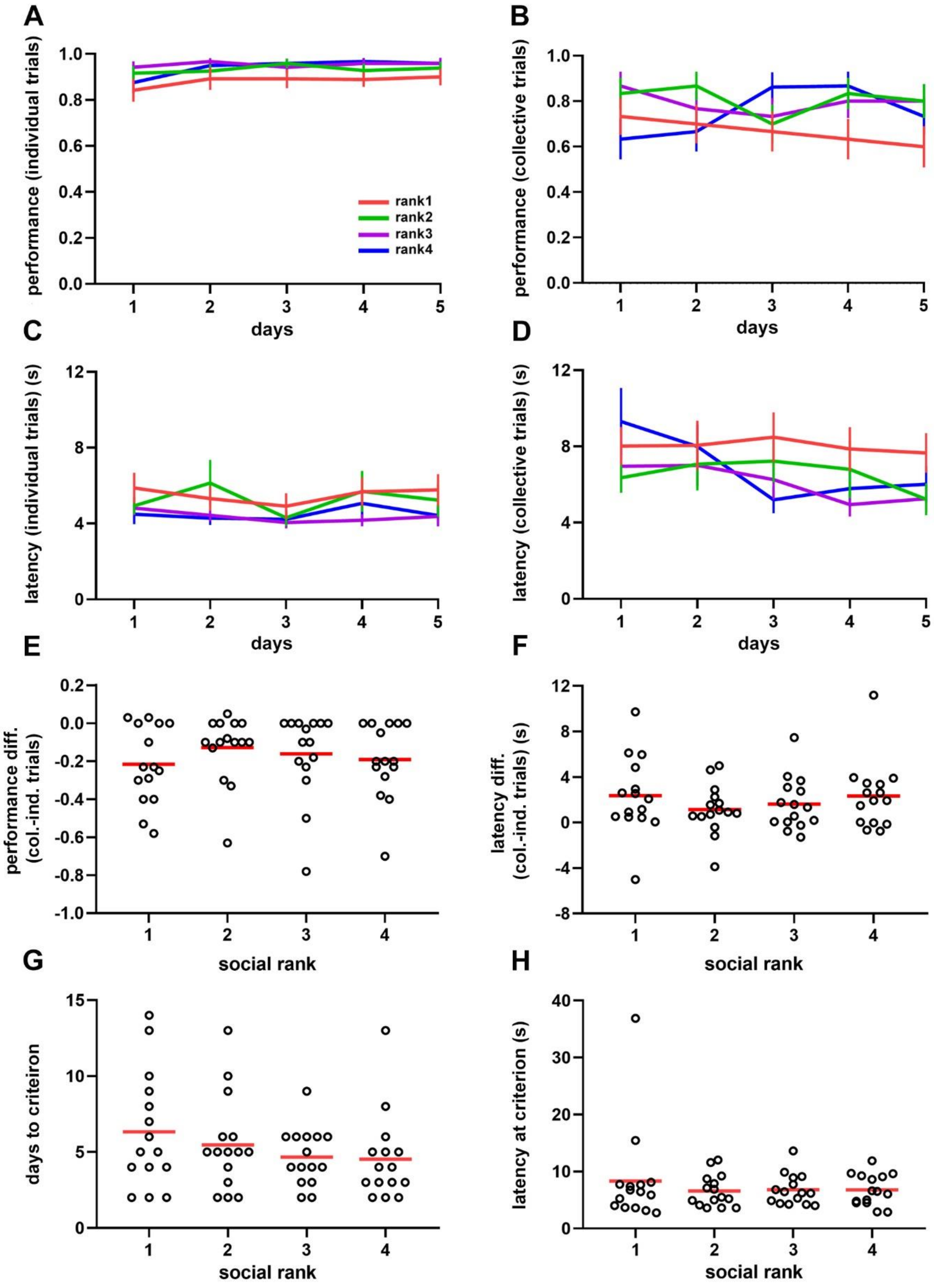

**Figure S7.** Average task performance (A, B) and latency (C, D) for individual (A, C) and collective (B, D) trials for all mice (n = 60) according to social ranking. Performance (E) and latency (F) difference between collective and individual trials during the testing phase of the task according to social ranking. One-way ANOVA, E, P = 0.655 F, P = 0.566. Time to reach learning criterion (G) and task latency at learning criterion (H) during the testing phase of the task according to social ranking. Kruskal-Wallis test, G, P = 0.4585; H, P = 0.9609. Colored lines, average population ± SEM; circles, average of individual animals; red lines; population average.

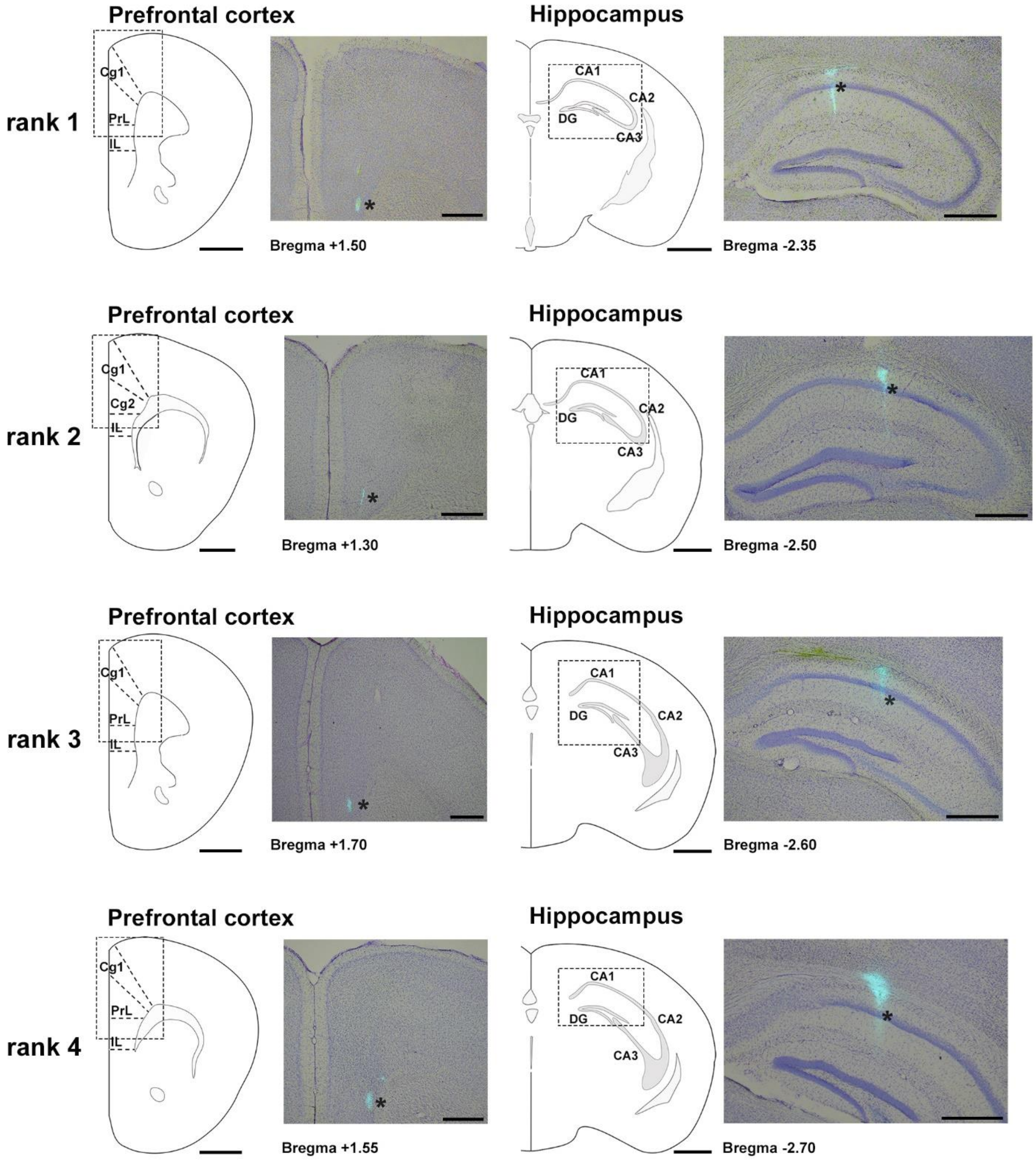

**Figure S8.** Anatomical location of recording electrodes. Examples sorted by social ranking. Ranking 1 (mouse CM99), ranking 2 (mouse CM65), ranking 3 (mouse CM64), ranking 4 (mouse CM47). Brain sections were Nissl stained and superimposed to fluorescent micrographs showing electrode tracks (blue, asterisks) in both cortex and hippocampus. Cg1, cingulate cortex; PrL, prelimbic cortex; IL, infralimbic cortex. CA1, CA2, CA3, cornu ammonis fields; DG, dentate gyrus. Scale bar 500 um.

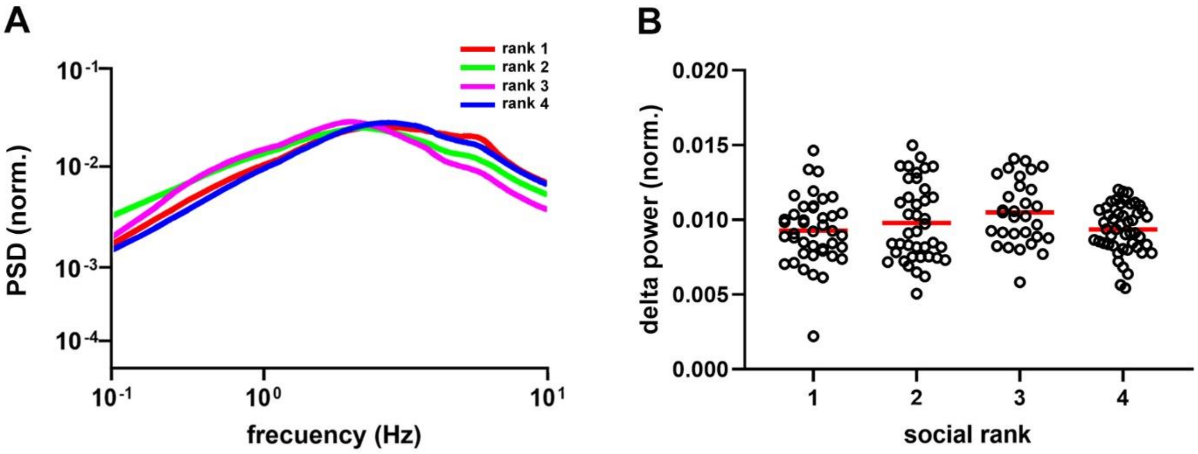

**Figure S9.** Cortical delta waves in anesthetized mice. Average (A) and peak (B) power spectral density of the prefrontal cortex of anesthetized mice according to social ranking (n = 22). One-way ANOVA, P = 0.081. Colored lines, averages; circles, record values; red lines; population average.

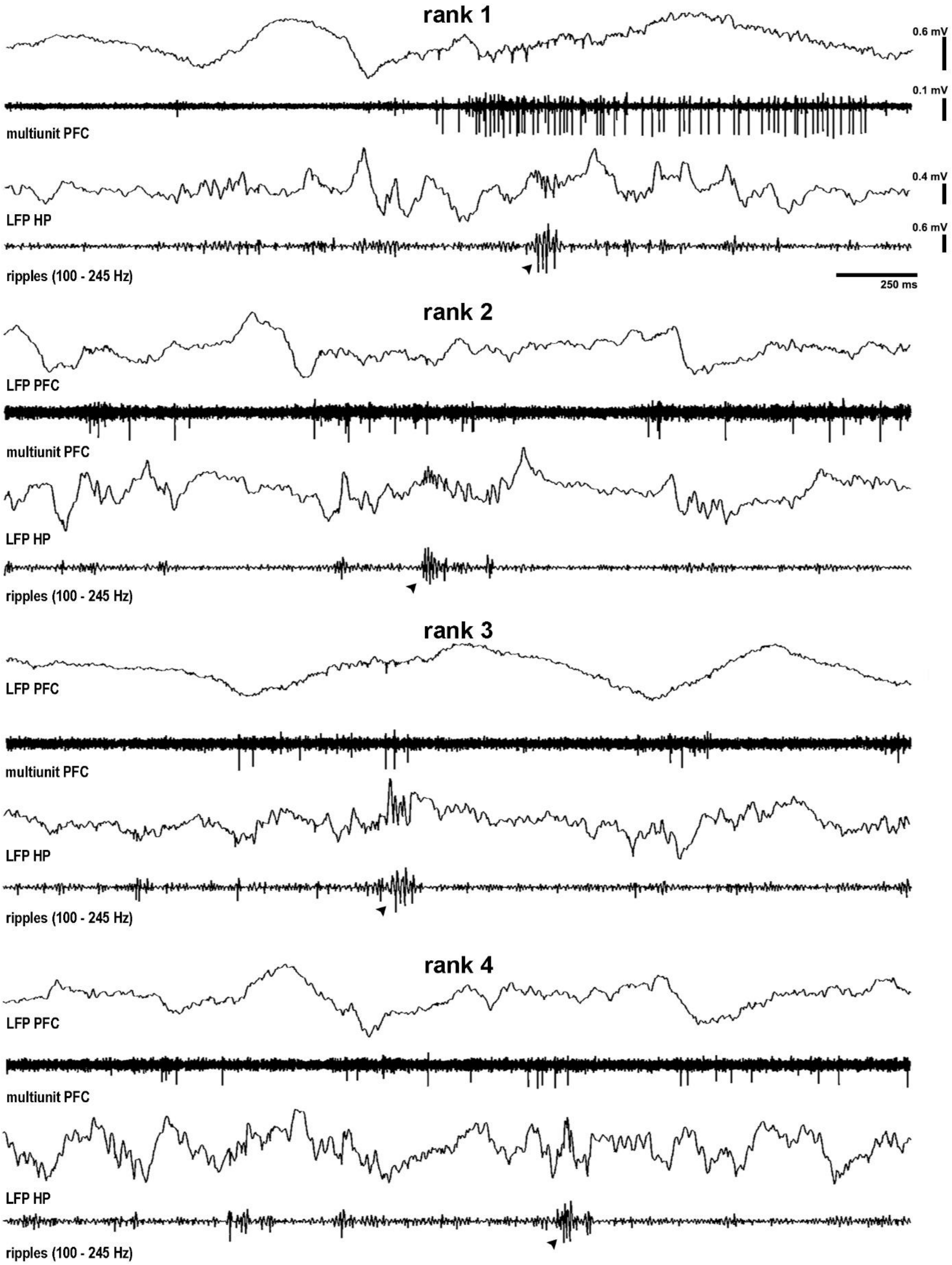

**Figure S10.** Examples of hippocampal sharp wave ripples and associated prefrontal cortex spiking for all social groups. Simultaneous recordings of prefrontal cortex (LFP PFC) and hippocampus (LFP HP) showing sharp wave ripples (ripples, filtered 100-250 Hz, arrowhead) and cortical spiking activity (multiunit PFC, filtered 300-4000 Hz) recorded in urethane-anesthetized mice: Rank 1, mouse CM24_reg05; rank 2, mouse CM73_reg02; rank 3, mouse CM28_reg02; rank 4, mouse CM98_reg01.

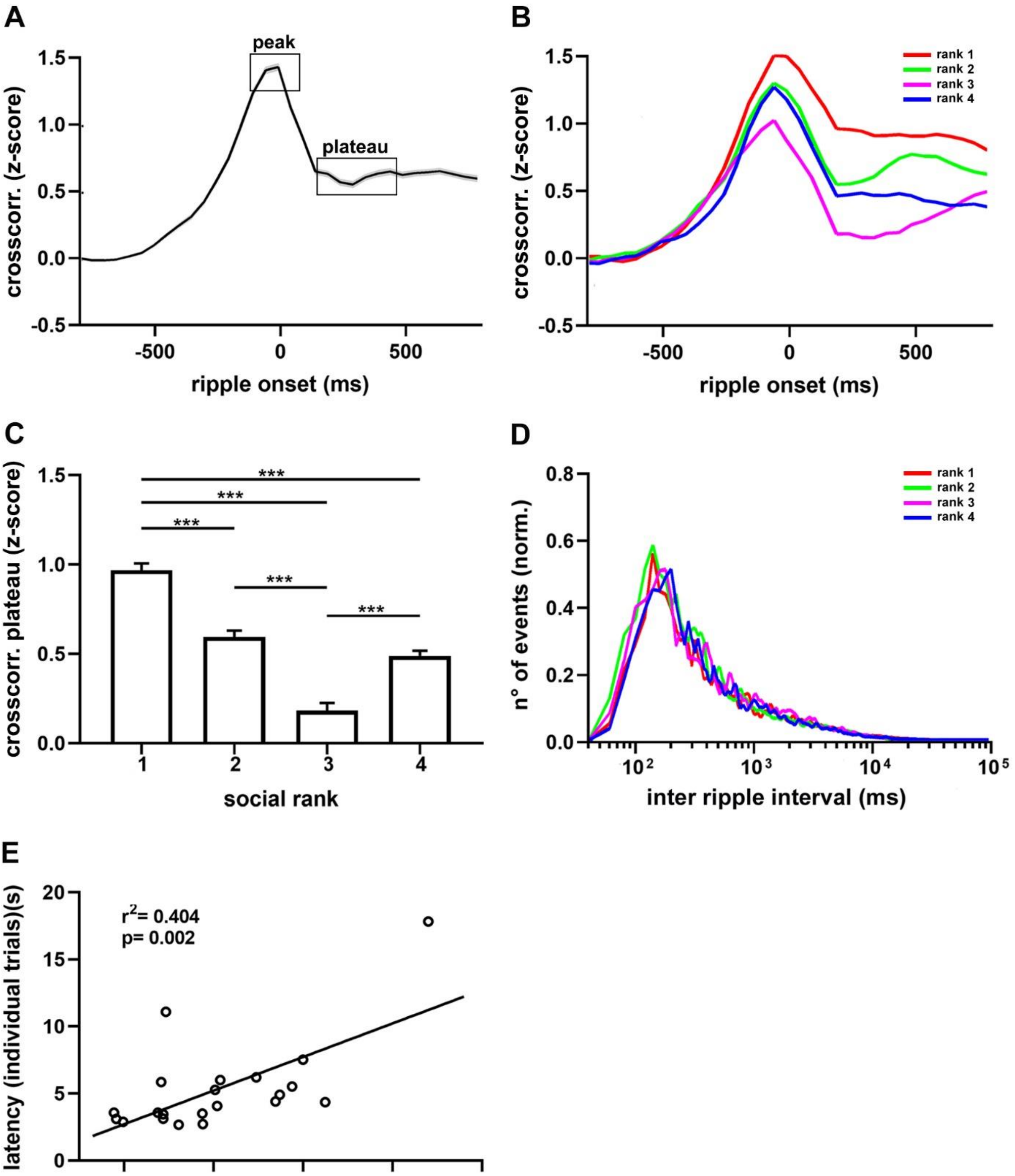

**Figure S11.** Functional connectivity between hippocampus and neocortex. A, average crosscorrelogram between the onset of sharp wave ripples and prefrontal units (n = 22 animals). B, average crosscorrelogram sorted by social ranking (n = 3702 units). C, crosscorrelogram plateau amplitude by social ranking. One-way ANOVA, P = 7.71e-47. D, Inter-event interval histogram for sharp wave ripples separated by social ranking. One-way ANOVA, P = 0.619. E, Spearman regression between crosscorrelogram's post-ripple plateau amplitude against task latency of individual trials. $R^2$ = 0.404, P = 0.0019. Bonferroni test post hoc (***, P < 0.001). Black lines, population averages; shading areas, ± SEM; colored lines, average population; bars, average ± SEM.

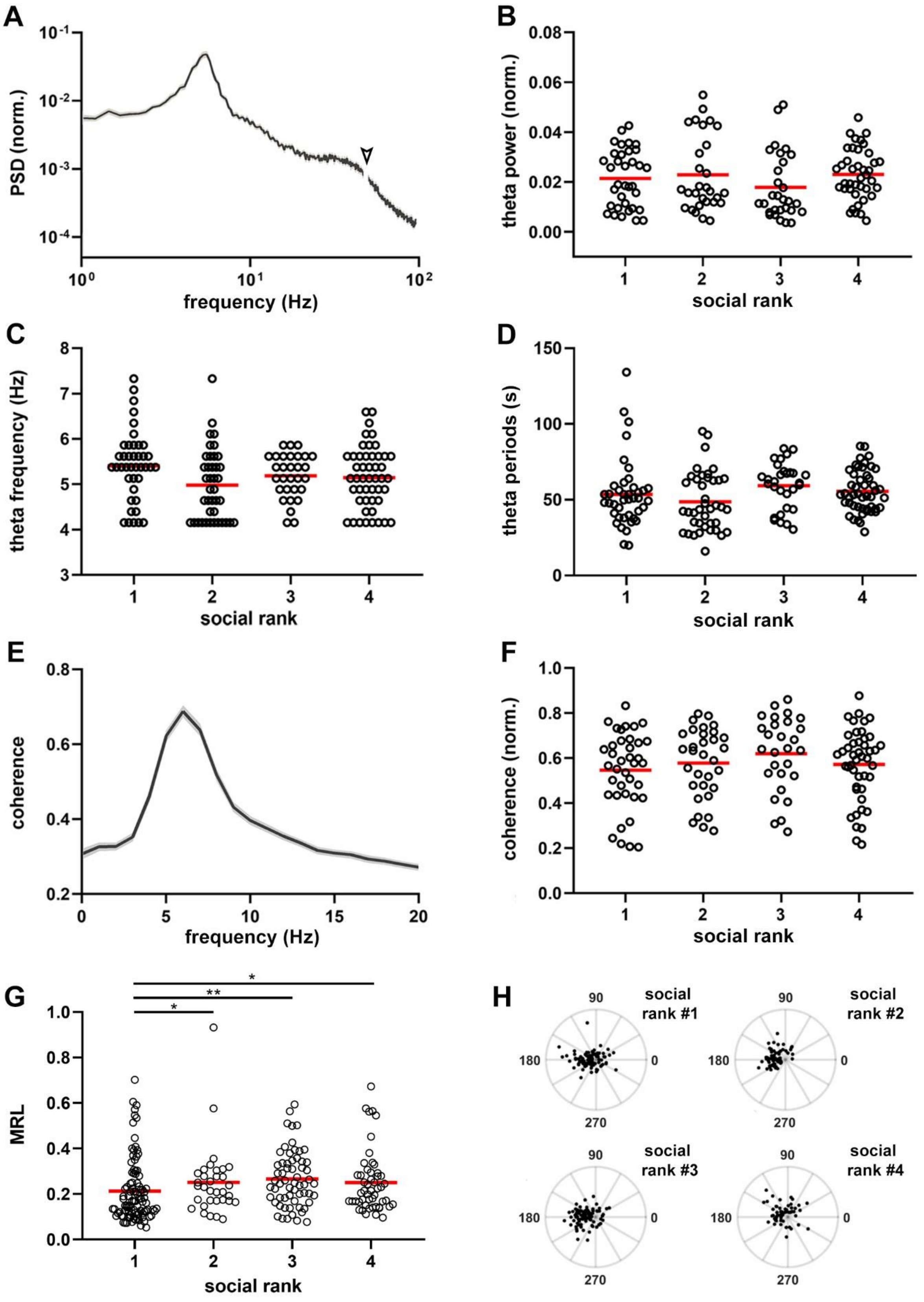

**Figure S12.** Theta oscillatory activity (4–8 Hz) in cortical networks. A, Average power spectral density (PSD) from all recorded animals (n = 21). Data are presented as mean ± SEM. B, Peak hippocampal theta amplitude by social ranking. One-way ANOVA, P = 0.3811. Peak frequency (C) and cumulative duration (D) of theta oscillations according to social ranking. One-way ANOVA; C, P = 0.0524; D, P = 0.0987. Average hippocampo-cortical spectral coherence (n = 21 animals). F, Peak hippocampo-cortical spectral coherence sorted by social ranking. One-way ANOVA, P = 0.350. G,

Average mean resultant length between prefrontal cortex spikes and hippocampal theta oscillations sorted by social ranking. Kruskal-Wallis test, $P = 0.0026$. H, Hippocampus theta phase modulation of cortical spikes sorted by social ranking. Each dot represents amplitude and angle of an individual unit. Red lines, population averages; circles, average of individual mice. Bonferroni test post hoc (*, $P < 0.05$; **, $P < 0.01$). Black lines, population averages; shading areas, ± SEM; circles, record values; red line; population average.

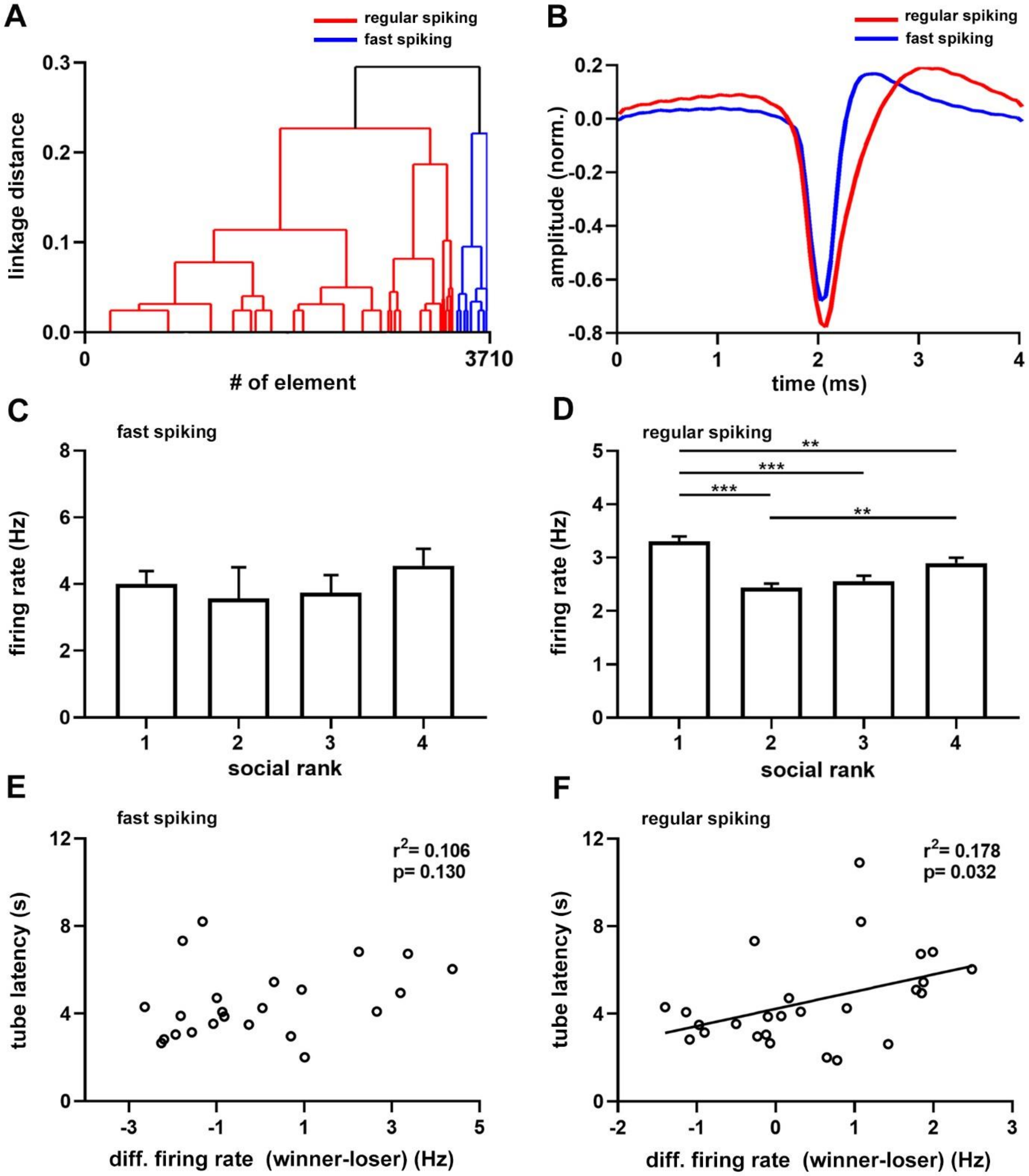

**Figure S13.** Cortical units recorded under anesthesia. A, dendrogram of all recorded units (n = 3702 units). Red shows largest cluster, regular spiking cells; blue shows smallest cluster, fast spiking cells. B, grand average of regular spiking (red, n = 3382) and fast spiking (blue, n = 320) units. Firing rate sorted by social ranking for fast spiking (C) and regular spiking (D) units. One-way ANOVA; C, P = 0.6243; D, P = 6.07e-12. Note that regular spiking cells in dominant animals discharge more than subordinate groups. Linear regression between the tube test latency difference (winner–loser) and the difference (winner–loser) of firing rate from regular spiking cells (E) or fast spiking cells (F). Only the linear regression for regular spiking cells was statistically significant. Bonferroni test post hoc (**, P < 0.01; ***, P < 0.001). Colored lines, average neuron waveforms; bars, average ± SEM.

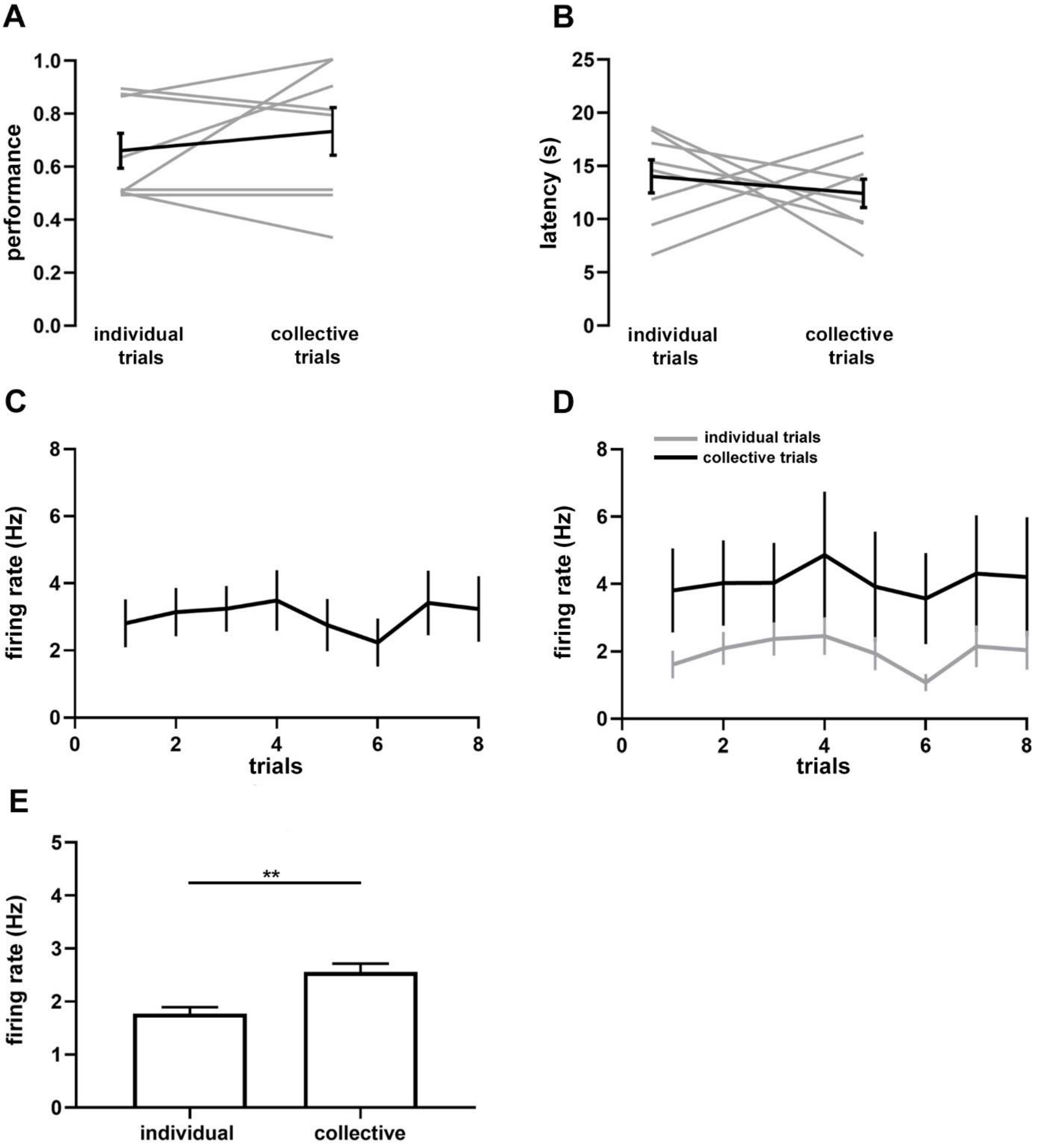

**Figure S14.** Average task performance (A) and latency (B) for chronically-implanted mice during collective and individual trials sampled during collective navigation (n = 8). One-sample t-test; A, P = 0.2597; B, P = 0.2217. C, average firing rate from all recorded units (n = 67) from chronically-implanted mice during task performance across trials. D, average firing rate from cortical units sorted by social context and trial number. E, average firing rate from cortical units during spatial navigation. Two-sample t-test, P = 0.000128. Two-sample t-test (**, P < 0.01). Gray lines, individual mice; black and gray lines, population averages ± SEM; bars, average ± SEM.

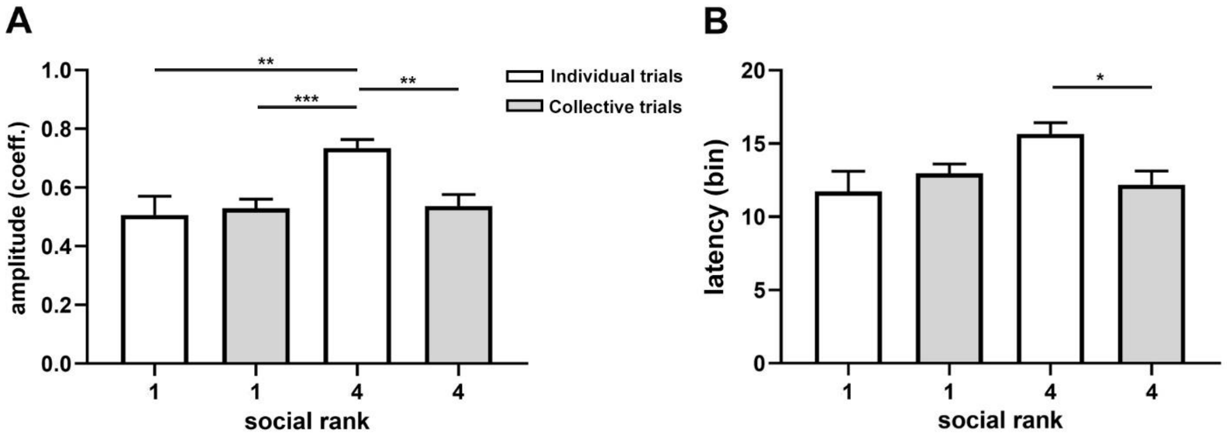

**Figure S15.** Variability in neuronal activity from the prefrontal cortex during spatial navigation. A, amplitude of spiking activity, calculated as the normalized firing rate from the last bins (15 to 20) subtracted from the first bins (1 to 5) divided by their total. Two-way ANOVA, P = 0.0103. B, latency of maximal spiking activity, calculated as the bin (1 to 20) at which each neuron discharged maximally. Two-way ANOVA, P = 0.0155. Bonferroni test post hoc (*, P < 0.05; **, P < 0.01; ***, P < 0.001). Bars, average ± SEM.

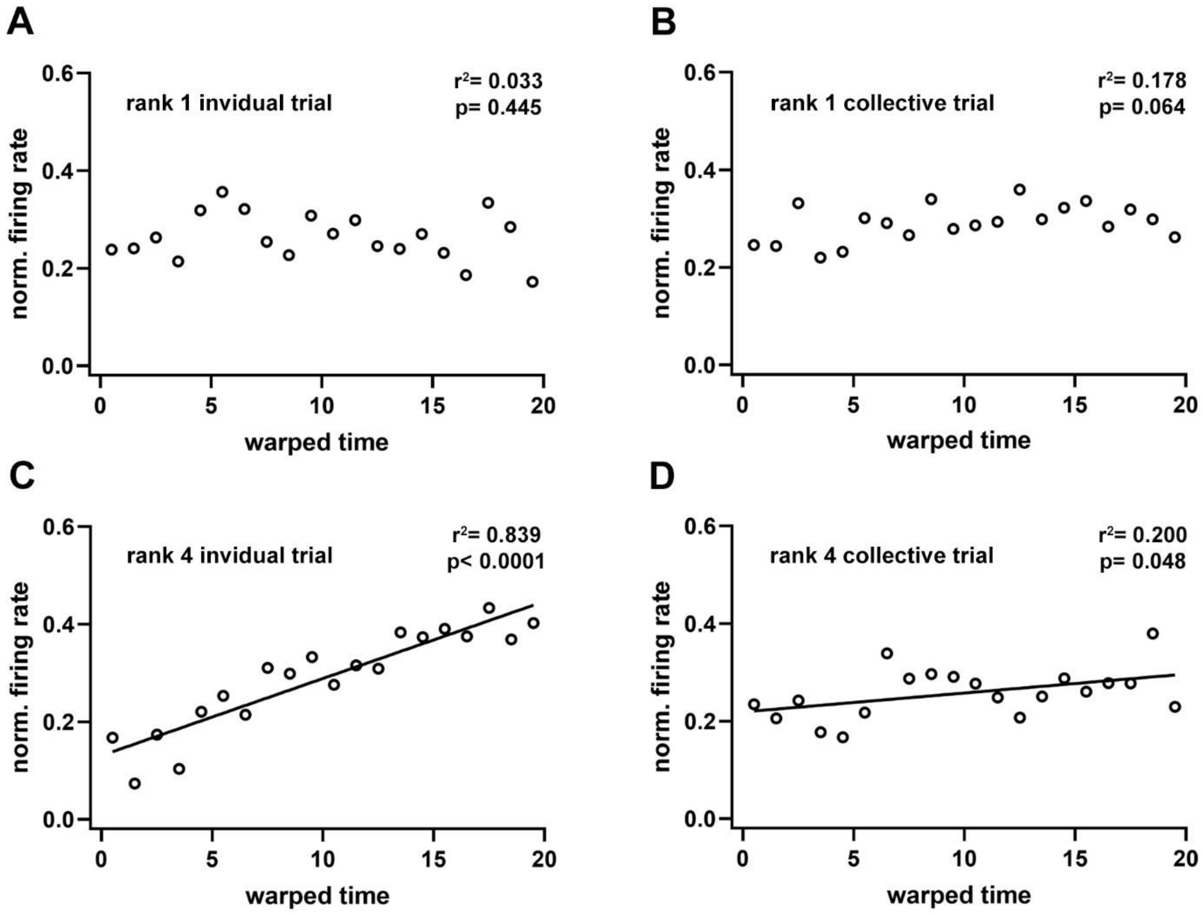

**Figure S16.** Normalized spiking activity in the prefrontal cortex during spatial navigation. Linear regressions of the normalized firing rates for dominant (A, B) and submissive (C, D) animals during T-maze navigation during individual (A, C) and collective (B, D) trials. Note that only submissive animals exhibit statistically significant regressions. Every point represents the average value from recorded neurons (dominant, n = 34; submissive, n = 33). Warped time represents entire T-maze. Bin 0, start box; bin 20, arm end.

### 4. SI tables

**Table S1.** P-values of linear regressions between pairs of neural and behavioral parameters

| | perf ind | perf coll | lat ind | lat coll | theta amp | theta fq | theta coh | ripples amp | ripples fq | cc peak | cc plateau | firing fq | pyr fq | int fq | PSI |
|---|---|---|---|---|---|---|---|---|---|---|---|---|---|---|---|
| **perf ind** | | e-5 | 0.01 | e-3 | 0.16 | 0.14 | 0.64 | 0.85 | 0.97 | 0.39 | 0.03 | 0.68 | 0.65 | 0.50 | 0.18 |
| **perf coll** | e-5 | | 0.49 | e-3 | 0.39 | 0.58 | 0.70 | 0.43 | 0.24 | 0.45 | 0.67 | 0.19 | 0.17 | 0.68 | 0.97 |
| **lat ind** | 0.01 | 0.49 | | e-5 | 0.05 | 0.01 | 0.99 | 0.94 | 0.93 | 0.58 | e-3 | 0.39 | 0.48 | e-3 | 0.07 |
| **lat coll** | e-3 | e-3 | e-5 | | 0.04 | 0.03 | 0.87 | 0.82 | 0.41 | 0.15 | 0.19 | 0.55 | 0.42 | 0.16 | 0.43 |
| **theta amp** | 0.16 | 0.39 | 0.05 | 0.04 | | 0.29 | 0.16 | 0.02 | 0.22 | 0.71 | 0.14 | 0.71 | 0.76 | 0.05 | 0.84 |
| **theta fq** | 0.14 | 0.58 | 0.01 | 0.03 | 0.29 | | 0.39 | 0.79 | 0.12 | 0.54 | e-3 | 0.93 | 0.75 | 0.29 | 0.08 |
| **theta coh** | 0.64 | 0.70 | 0.99 | 0.87 | 0.16 | 0.39 | | 0.59 | 0.03 | 0.86 | 0.63 | 0.53 | 0.81 | 0.06 | 0.74 |
| **ripples amp** | 0.85 | 0.43 | 0.94 | 0.82 | 0.02 | 0.79 | 0.59 | | 0.03 | 0.77 | 0.67 | 0.82 | 0.42 | 0.82 | 0.62 |
| **ripples fq** | 0.97 | 0.24 | 0.93 | 0.41 | 0.22 | 0.12 | 0.03 | 0.03 | | 0.46 | 0.95 | 0.89 | 0.37 | 0.78 | 0.92 |
| **cc peak** | 0.39 | 0.45 | 0.58 | 0.15 | 0.71 | 0.54 | 0.86 | 0.77 | 0.46 | | 0.24 | 0.13 | 0.10 | 0.51 | 0.87 |
| **cc plateau** | 0.03 | 0.67 | e-3 | 0.19 | 0.14 | e-3 | 0.63 | 0.67 | 0.95 | 0.24 | | 0.57 | 0.68 | 0.09 | e-3 |
| **firing fq** | 0.68 | 0.19 | 0.39 | 0.55 | 0.71 | 0.93 | 0.53 | 0.82 | 0.89 | 0.13 | 0.57 | | e-15 | e-3 | 0.08 |
| **pyr fq** | 0.65 | 0.17 | 0.48 | 0.42 | 0.76 | 0.75 | 0.81 | 0.42 | 0.37 | 0.10 | 0.68 | e-15 | | 0.01 | 0.78 |
| **int fq** | 0.50 | 0.68 | e-3 | 0.16 | 0.05 | 0.29 | 0.06 | 0.82 | 0.78 | 0.51 | 0.09 | e-3 | 0.01 | | 0.09 |
| **PSI** | 0.18 | 0.97 | 0.07 | 0.43 | 0.84 | 0.08 | 0.74 | 0.62 | 0.92 | 0.87 | e-3 | 0.08 | 0.78 | 0.09 | |

Performance in individual trials, perf ind; performance in collective trials, perf coll; latency in individual trials, lat ind; latency in collective trials, lat coll; theta oscillation amplitude, theta amp; theta oscillation frequency, theta fq; theta oscillation coherence, theta coh; sharp wave ripples amplitude, ripple amp; sharp wave ripples frequency, ripple fq; crosscorrelogram peak amplitude, cc peak; crosscorrelogram plateau amplitude, cc plateau; neuronal firing rate; firing fq; pyramidal neurons firing rate, pyr fq; interneurons firing rate; int fq; peer sensitivity index, PSI. 10e-3, e-3; 10e-5, e-5; 10e-15, e-15. Yellow squares show uncorrected significant correlations, $P < 0.05$.

**Table S2.** Mixed logistic model with fixed effects for collective task performance (n = 60 animals used for behavioral tests)

| Parameter | estimate | SE | P |
|---|---|---|---|
| intercept | 0.2721 | 0.8258 | 0.741770 |
| individual performance | 2.8035 | 0.8182 | *0.000612* |
| animal density (selected arm) | -3.1858 | 0.4876 | *6.41e-11* |
| animal density (opposite arm) | -1.5323 | 0.4978 | *0.002085* |
| total animal density | 1.0492 | 0.3206 | *0.001066* |

**Table S3.** Mixed logistic model with fixed effects for collective task performance (n = 20 animals used for anesthesia recordings)

| **Parameter** | **estimate** | **SE** | **P** |
|---|---|---|---|
| intercept | -0.1794 | 1.3608 | 0.89513 |
| individual performance | 3.1845 | 1.3582 | *0.01905* |
| animal density (selected arm) | -2.4680 | -3.075 | *0.00211* |
| animal density (opposite arm) | -1.1839 | 0.7862 | 0.13 |
| total animal density | 0.78 | 0.52 | 0.13 |

**Table S4.** Multiple linear model for the PSI (n = 20 animals used for anesthesia recordings)

| **Parameter** | **estimate** | **SE** | **P** |
|---|---|---|---|
| intercept | 58.76 | 35.37 | 0.1206 |
| neuronal firing rate | -23.24 | 12.23 | 0.0798 |
| mean resultant length | 40.08 | 142.05 | 0.7823 |
| crosscorrelogram ripple-peak | 11.86 | 10.72 | 0.2888 |
| crosscorrelogram post-ripple | -26.43 | 10.37 | *0.0242* |

**Table S5.** Multiple linear model for the PSI (n = 20 animals used for anesthesia recordings)

| **Parameter** | **estimate** | **SE** | **P** |
|---|---|---|---|
| intercept | -9.7985 | 151.0599 | 0.949 |
| ripple frequency | 0.1181 | 1.0649 | 0.913 |
| ripple amplitude | 0.4150 | 18.5729 | 0.982 |
| crosscorrelogram ripple-peak | 5.9279 | 11.9300 | 0.626 |
| crosscorrelogram post-ripple | 28.1367 | 10.0703 | *0.013* |

**Table S6.** Multiple linear model for individual task performance (n = 20 animals used for anesthesia recordings)

| **Parameter** | **estimate** | **SE** | **P** |
|---|---|---|---|
| intercept | 0.843824 | 0.185007 | *0.000375* |
| collective performance | 0.346942 | 0.046636 | *2.08e-06* |
| ripple frequency | -0.002180 | 0.001444 | 0.151925 |
| ripple amplitude | 0.039210 | 0.025525 | 0.145326 |
| crosscorrelogram peak-ripple | 0.005932 | 0.016553 | 0.725076 |
| crosscorrelogram post-ripple | -0.050519 | 0.012887 | *0.001364* |

**Table S7.** Multiple linear model for individual task latency (n = 20 animals used for anesthesia recordings)

| **Parameter** | **estimate** | **SE** | **P** |
|---|---|---|---|
| intercept | -0.14242 | 9.83315 | 0.98863 |
| collective latency | 0.47684 | 0.15791 | *0.00862* |
| ripple frequency | 0.01737 | 0.07250 | 0.81386 |
| ripple amplitude | -0.48678 | 1.20544 | 0.69204 |
| crosscorrelogram peak-ripple | -0.04911 | 0.90651 | 0.95751 |
| crosscorrelogram post-ripple | 1.79839 | 0.69635 | *0.02081* |

**Table S8.** Univariate tests of significance for firing rate (n = 20 animals used for anesthesia recordings)

| **parameter** | **SS** | **F** | **P** |
|---|---|---|---|
| intercept | 10187.48 | 1169.875 | *10e-6* |
| cortical region | 15.66 | 1.799 | 0.179932 |
| neuron type | 325.64 | 37.395 | *10e-6* |
| hierarchy | 93.78 | 3.590 | *0.013127* |
| cortical region*neuron type | 20.00 | 2.297 | 0.129738 |
| cortical region*hierarchy | 64.68 | 2.476 | 0.059632 |
| neuron type*hierarchy | 40.71 | 1.558 | 0.197450 |
| cortical region*neuron type*hierarchy | 64.79 | 2.480 | 0.059302 |

SS, sum of squares; F, F-statistic; P, p-value; cortical region (dorsal or ventral prefrontal cortex); neuron type (fast spiking or regular spiking cells); hierarchy (dominant, first active subordinate, second active subordinate, or submissive).

**Table S9.** Multiple linear model for latency difference (winner – loser) in the tube test (n = 20 animals used for anesthesia recordings)

| **Parameter** | **estimate** | **SE** | **P** |
|---|---|---|---|
| intercept | 4.1093 | 0.5823 | *2.11e-05* |
| neuronal firing rate (difference) | 1.1343 | 0.3552 | *0.00855* |
| mean resultant length (difference) | 2.7736 | 9.3354 | 0.77192 |
| crosscorrelogram ripple-peak (difference) | -0.6268 | 0.5366 | 0.26739 |
| crosscorrelogram post-ripple (difference) | 0.3177 | 0.4178 | 0.46298 |

**Table S10.** Multiple linear model for individual task performance (n = 20 animals used for anesthesia recordings)

| **Parameter** | **estimate** | **SE** | **P** |
|---|---|---|---|
| intercept | 1.05821 | 0.24175 | *0.000469* |
| collective performance | 0.31042 | 0.05827 | *6.8e-05* |
| theta power | -1.62367 | 1.76202 | 0.370484 |
| theta frequency | -0.06966 | 0.04583 | 0.148007 |
| theta coherence | 0.04803 | 0.12314 | 0.701632 |

**Table S11.** Multiple linear model for individual task latency (n = 20 animals used for anesthesia recordings)

| **Parameter** | **estimate** | **SE** | **P** |
|---|---|---|---|
| intercept | -11.6517 | 9.2605 | 0.22637 |
| collective latency | 0.4623 | 0.1547 | *0.00871* |
| theta power | 29.9151 | 70.4885 | 0.67693 |
| theta frequency | 2.2370 | 1.8760 | 0.25048 |
| theta coherence | 2.6051 | 4.5705 | 0.57660 |

**Table S12.** Univariate tests of significance for firing rate (n = 8 animals used for chronic recordings)

| **parameter** | **SS** | **F** | **P** |
|---|---|---|---|
| intercept | 4241.786 | 464.746 | *10e-6* |
| state | 77.605 | 8.503 | *0.003631* |
| hierarchy | 315.523 | 34.569 | *10e-6* |
| context | 251.525 | 27.556 | *10e-6* |
| state*hierarchy | 94.639 | 10.369 | *0.001326* |
| state*context | 15.048 | 1.649 | 0.199447 |
| hierarchy*context | 0.018 | 0.0019 | 0.965011 |
| state*hierarchy*context | 23.310 | 2.5539 | 0.110361 |

SS, sum of squares; F, F-statistic; P, p-value; hierarchy, dominance hierarchy (dominant or submissive); state, behavioral state (home cage or spatial navigation); context, social context (individual or collective trial).

**Table S13.** Univariate tests of significance for mean resultant length (n = 8 animals used for chronic recordings)

| **parameter** | **SS** | **F** | **P** |
|---|---|---|---|
| intercept | 1.932116 | 106.5667 | *10e-6* |
| state | 0.100625 | 5.55 | *0.020049* |
| hierarchy | 0.198114 | 10.9271 | *0.00124* |
| context | 0.001088 | 0.06 | 0.806913 |
| state*hierarchy | 0.204661 | 11.2882 | *0.001037* |
| state*context | 0.026326 | 1.452 | 0.230494 |
| hierarchy*context | 0.014867 | 0.82 | 0.36694 |
| state*hierarchy*context | 0.000219 | 0.0121 | 0.912703 |

SS, sum of squares; F, F-statistic; p, p-value; hierarchy, dominance hierarchy (dominant or submissive); state, behavioral state (home cage or spatial navigation); context, social context (individual or collective trial).

**Table S14.** Univariate tests of significance for amplitude of maximal spiking activity (n = 8 animals used for chronic recordings)

| **parameter** | **SS** | **F** | **p** |
|---|---|---|---|
| Intercept | 57.93621 | 730.1778 | *0.000000* |
| hierarchy | 0.60437 | 7.6169 | *0.006259* |
| context | 0.33079 | 4.1690 | *0.042336* |
| hierarchy *context | 0.53184 | 6.7028 | *0.010253* |

SS, sum of squares; F, F-statistic; P, p-value; hierarchy, dominance hierarchy (dominant or submissive); context, social context (individual or collective trial).

**Table S15.** Univariate tests of significance for latency of maximal spiking activity (n = 8 animals used for chronic recordings)

| **parameter** | **SS** | **F** | **p** |
|---|---|---|---|
| Intercept | 30140.54 | 742.6998 | *0.000000* |
| hierarchy | 108.81 | 2.6812 | 0.102932 |
| context | 52.95 | 1.3048 | 0.254546 |
| hierarchy *context | 241.54 | 5.9518 | *0.015473* |

SS, sum of squares; F, F-statistic; p, p-value; hierarchy, dominance hierarchy (dominant or submissive); context, social context (individual or collective trial).

**Table S16.** Summary of statistical tests per figure

| Figure | Test | Condition | Statistic | P |
|---|---|---|---|---|
| 1B | Wilcoxon signed rank test | performance | Zval = 5.5584 | 2.72e-08 |
| 1C | Wilcoxon signed rank test | latency | Zval = -5.0317 | 4.86e-07 |
| 1D | Spearman correlation | individual performance/collective performance | r2 = 0.501 | 1.5e-10 |
| 1E | Spearman correlation | animal density/collective performance | r2 = 0.220 | 1.6e-04 |
| 2B | one-way ANOVA | time in tube | F(5,84) = 20.58 | 2.31e-13 |
| 2C | one-way ANOVA | individual performance | F(3,56) = 1.41 | 0.25 |
| 2D | one-way ANOVA | animal density (selected arm) | F(3,580) = 4.11 | 0.007 |
| 2E | one-way ANOVA | PSI | F(3,56) = 4.3 | 0.0084 |
| 3C | one-way ANOVA | ripple amplitude | F(3,18567) = 27.59 | 8.55e-18 |
| 3D | one-way ANOVA | ripple frequency | F(3,18567) = 59.06 | 5.45e-38 |
| 3E | one-way ANOVA | duration of ripples | F(3,18567) = 2.4 | 0.0658 |
| 3F | one-way ANOVA | amplitude (peak crosscorr.) | F(3,654) = 4.92 | 0.002 |
| 3G | one-way ANOVA | amplitude (plateau crosscorr.) | F(3,654) = 11.37 | 2.81e-07 |
| 3H | Pearson correlation | PSI/amplitude | r2 = 0.328 | 0.00666 |
| 3I | Pearson correlation | performance/amplitude | r2 = 0.235 | 0.026 |
| 4B | one-way ANOVA | firing rate | F(3,3698) = 17.03 | 5.56e-11 |
| 4C | Kruskal-Wallis test | MRL theta PFC (anesthesia) | Chi-sq = 28.33 | 3.10e-06 |
| 4D | Pearson correlation | latency (time in interaction tube)/diff firing rate | r2 = 0.226 | 0.0142 |
| 5B | two-way ANOVA | firing rate (homecage & navigation) | F(7,929) = 10.37 | 0.001326 |
| 5C | two-way ANOVA | MRL (homecage &navigation) | F (7,124) = 11.29 | 0.001037 |
| 5D | Wilcoxon rank sum test | norm firing rate rank#1 (warped) | | < 0.05 |
| 5E | Wilcoxon rank sum test | norm firing rate rank#4 (warped) | | < 0.05 |
| S3E | Spearman correlation | latency (collective)/latency (individual) | r2 = 0.529 | 4.4e-11 |

| | | | | |
|---|---|---|---|---|
| S3F | Pearson correlation | performance (collective)/performance (individual) | r2 = 0.642 | 1.45e-14 |
| S4A | Spearman correlation | performance (collective)/density | r2 = 0.019 | 0.293 |
| S4B | one-way ANOVA | animal density (opposite arm) | F(3,56) = 0.89 | 0.4499 |
| S4C | one-way ANOVA | animal density (opposite+ selected arm) | F(3,56) = 1.7 | 0.1783 |
| S6E | one-way ANOVA | initial body mass | F(3,56) = 0.31 | 0.8216 |
| S6F | one-way ANOVA | final body mass | F(3,56) = 0.54 | 0.6554 |
| S6G | one-way ANOVA | % of body weight change | F(3,56) = 0.01 | 0.983 |
| S7E | one-way ANOVA | performance difference | F(3,56) = 0.54 | 0.6552 |
| S7F | one-way ANOVA | latency difference | F(3,56) = 0.68 | 0.5662 |
| S7G | Kruskal-Wallis test | days to criterion | Chi-sq = 2.59 | 0.4585 |
| S7H | Kruskal-Wallis test | latency at criterion | Chi-sq = 0.3 | 0.9609 |
| S9B | one-way ANOVA | delta power | F(3,155) = 2.29 | 0.081 |
| S11C | one-way ANOVA | crosscrorr. plateau | F(3,3698) = 74.59 | 7.71e-47 |
| S11D | one-way ANOVA | inter ripple interval | F(3,19992) = 0.59 | 0.6193 |
| S11E | Pearson correlation | latency/amplitude | r2 = 0.404 | 0.0019 |
| S12B | one-way ANOVA | theta power | F(3,131) = 1.03 | 0.3811 |
| S12C | one-way ANOVA | theta frequency | F(3,155) = 2.63 | 0.0524 |
| S12D | one-way ANOVA | theta periods | F(3,155) = 2.13 | 0.0987 |
| S12F | one-way ANOVA | theta coherence | F(3,139) = 1.1 | 0.35 |
| S12G | Kruskal-Wallis test | MRL theta hippocampus (anesthesia) | Chi-sq = 14.27 | 0.0026 |
| S13C | one-way ANOVA | rate (fast spiking) | F(3,316) = 0.59 | 0.6243 |
| S13D | one-way ANOVA | rate (regular spiking) | F(3,3378) = 18.57 | 6.07e-12 |
| S13E | Pearson correlation | latency (time in tube)/diff. firing rate - fast spiking rate | r2 = 0.106 | 0.13 |
| S13F | Pearson correlation | latency/diff. firing rate - regular spiking rate | r2 = 0.178 | 0.032 |
| S14A | one-sample t-test | performance | tstat = -1.2244 | 0.2597 |
| S14B | one-sample t-test | latency | tstat = 1.3412 | 0.2217 |
| S14E | two-sample t-test | firing rate/testing phase | tstat = -3.8465 | 0.000128 |
| S15A | two-way ANOVA | norm. firing rate during spatial navigation | F(3,225) = 6.70 | 0.010253 |
| S15B | two-way ANOVA | bin max during spatial navigation | F(3,226) = 5.95 | 0.015473 |
| S16A | Pearson correlation | norm. firing rate individual trial rank #1 | r2 = 0.0327 | 0.4454 |
| S16B | Pearson correlation | norm. firing rate collective trial rank #1 | r2 = 0.1779 | 0.0640 |

| | | | | |
|---|---|---|---|---|
| S16C | Pearson correlation | norm. firing rate individual trial rank #4 | r2 = 0.8394 | 0.0000143 |
| S16D | Pearson correlation | norm. firing rate collective trial rank #4 | r2 = 0.2000 | 0.0481 |

### 5. SI Videos

| video | file | trial | mouse | cage | description |
|---|---|---|---|---|---|
| S1 | tube_test | 5 | ti01, ca02 | 212A | Two littermates in the tube test |
| S2 | individual_acute | 1 | ti1844 | 339A | T-maze individual trial of dominant mouse |
| S3 | collective_acute | 3 | ti1844, td185, cm1846, cn1847 | 339A | T-maze collective trial of four littermates |
| S4 | individual_chronic | 6 | H8372 | N/A | T-maze individual trial of implanted dominant mouse |
| S5 | collective_chronic | 3 | H8372, H8373 | N/A | T-maze collective trial of two implanted littermates |
